# A Universal Allosteric Mechanism for G Protein Activation

**DOI:** 10.1101/2020.11.15.383679

**Authors:** Kevin M. Knight, Soumadwip Ghosh, Sharon L. Campbell, Tyler J. Lefevre, Reid H. J. Olsen, Alan V. Smrcka, Natalie H. Valentin, Guowei Yin, Nagarajan Vaidehi, Henrik G. Dohlman

**Affiliations:** Department of Pharmacology, University of North Carolina at Chapel Hill, Chapel Hill, NC 27599, USA; Department of Computational and Quantitative Medicine, Beckman Research Institute of the City of Hope, Duarte, CA 91010, USA; Department of Biochemistry and Biophysics, University of North Carolina at Chapel Hill, Chapel Hill, NC 27599, USA; Department of Pharmacology, University of Michigan, Ann Arbor, MI 48109, USA

**Author notes:** Corresponding authors: Henrik G. Dohlman, Nagarajan Vaidehi.

**Keywords:** G protein, molecular dynamics simulations, nuclear magnetic resonance spectroscopy, bioluminescence resonance energy transfer

## Abstract

G proteins play a central role in signal transduction and pharmacology. Signaling is initiated by cell-surface receptors, which promote GTP binding and the dissociation of Gα from the Gβγ subunits. Structural studies have revealed the molecular basis for subunit association with receptors, RGS proteins and downstream effectors. In contrast, the mechanism of subunit dissociation is poorly understood. We used cell signaling assays, MD simulations, biochemistry and structural analysis to identify a conserved network of amino acids that dictates subunit release. In the presence of the terminal phosphate of GTP, a glycine forms a polar network with an arginine and glutamate, putting torsional strain on the subunit binding interface. This “G-R-E motif” secures GTP and, through an allosteric link, discharges the Gβγ dimer. Replacement of network residues prevents subunit dissociation, regardless of agonist or GTP binding. These findings reveal the molecular basis for the final committed step of G protein activation.

**HIGHLIGHTS:**

- Receptors promote GTP-GDP exchange and dissociation of G protein α and βγ subunits
- We find an allosteric network linking the γ phosphate of GTP with release of Gβγ
- The network consists of a conserved Gly-Arg-Glu “activation triad”
- Triad mutations prevent subunit dissociation, regardless of agonist or GTP binding
- Triad mutations are responsible for human endocrine and neurological disorders

## INTRODUCTION

Many hormones, neurotransmitters, and clinically important drugs elicit their effects through G protein coupled receptors (GPCRs). Receptors of this class represent one of the largest gene families, and by far the largest class of drug targets. Upon receptor activation by agonist, the G protein α subunit releases GDP, binds GTP, and dissociates from the Gβγ dimer. The free Gα subunit then modulates the activity of downstream effector proteins such as adenylyl cyclase and phospholipase C, among others. The Gβγ dimer can regulate many of these same effectors as well as potassium channels, calcium channels, phosphatidylinositol-3 kinase and MAPKs. Signaling is terminated after Gα hydrolyzes GTP and the heterotrimer reassembles. In most cases this inactivation step is accelerated by RGS proteins. Thus, receptors serve as signal discriminators while RGS proteins serve as timing devices and G proteins serve as determinants of output specificity.

The effects of G protein activation -- such as vasodilation, pain reduction, euphoria as well as light and odor sensing -- are quite diverse depending on the cellular context. Additional diversity comes from differential expression of 800+ GPCRs, which includes members with little or no shared sequence similarity. In contrast, G proteins are highly conserved in structure and function, and in mammals consist of only four subfamilies (G_s_, G_i/o_, G_q/11_ and G_12/13_) encoded by 16 individual genes.

The ability of such widely divergent receptors to activate a comparatively small number of G proteins indicates a shared mechanism of activation (Flock et al., 2017; Flock et al., 2015). Broadly speaking, this process can be broken down into three discrete protein transformations: (i) ligand-specific conformational changes in the receptor, (ii) GTP-induced conformational changes in Gα, and (iii) displacement of the Gβγ subunits. It had long been postulated that an activated receptor must induce an opening of the two domains within Gα, stabilizing the nucleotide free state and facilitating GTP-GDP exchange. Direct evidence of such inter-domain opening was first provided through double electron-electron resonance experiments on rhodopsin-activated Gα_i_ (Van Eps et al., 2011) and on Gα_s_ (Dror et al., 2015), hydrogendeuterium-exchange studies (Chung et al., 2011) and through X-ray crystallographic studies of the β_2_ adrenergic receptor bound to the G_s_ protein heterotrimer (Rasmussen et al., 2011). Collectively, these findings indicate that Gα undergoes a “clam-shell like’ opening to facilitate nucleotide release, and that both domains contribute to the process. In all of these structures however, Gβγ remains associated with Gα, even in the nucleotide-free and agonist-bound complex. Thus, the conformational changes associated with the release of nucleotide are necessary but not sufficient for the subsequent release of Gβγ.

The third and final step of G protein activation begins with GTP binding. This entails conformational changes in three segments of Gα known as Switch I, II and III (Lambright et al., 1994). Crystal structures revealed that that Switch I connects the two domains of the Gα subunit, comprised of a RAS-homologous domain and an all-helical domain. Switches II and III are located within the RAS-like domain and are disordered in the GDP-bound structure. Upon GTP binding Switch II forms a well-ordered helix and makes multiple contacts with residues in the α3 helix. Switches I and II interact directly with the γ-phosphate of GTP and form part of the Gβγ-interacting surface (Lambright et al., 1996; Wall et al., 1995). Therefore, to form the active state, residues critical to maintaining Gβγ binding must retract inward toward the newly bound GTP molecule, creating a “molecular tug-of-war.” Less is known about the mechanism behind the dynamic cascade of events linking the γ-phosphate to Switch I and II and the displacement of Gβγ. Here, by combining cross-species sequence analysis, Molecular Dynamics (MD) simulations, thermostability measurements, direct Gα and Gβγ binding assays, NMR chemical shift analysis and cell-based functional assays we have uncovered the dynamic and transient effect of GTP binding leading to Gβγ dissociation. These models were also tested functionally, in biological contexts including the model organism *S. cerevisiae*. Our analysis reveals the existence of an evolutionarily conserved Gly-Arg-Glu triad that underpins all G protein activation. In particular, we show that these residues form an allosteric link between GTP and Gβγ. Mutations in any of these residues impair subunit dissociation, most likely due to a loss of a polar network between the terminal (γ) phosphate of GTP and the Gβγ-binding interface. Given that this motif is perfectly conserved among known Gα subunits, it is likely to be a universal feature of G protein activity. Thus, our findings reveal how a single phosphate group controls G protein subunit dissociation and signaling, processes that underlie much of human physiology and a substantial fraction of medical pharmacology.

## RESULTS

### Triad hypothesis

The active state of a receptor is able to bind simultaneously to agonist and nucleotide-free G protein; agonist binding stabilizes the empty form of the G protein heterotrimer, thereby facilitating the exchange of GTP for GDP. Binding to either nucleotide favors release of G protein from receptor and of receptor from agonist (De Lean et al., 1980; DeVree et al., 2016). However, it is GTP alone that triggers the release of Gβγ from Gα (Figure 1). Thus, the active state of the G protein can be defined as the conformation stabilized by the terminal phosphate of GTP. Our goal here was to determine the molecular rearrangements responsible for this third and final committed step of G protein activation.

**Figure 1.**
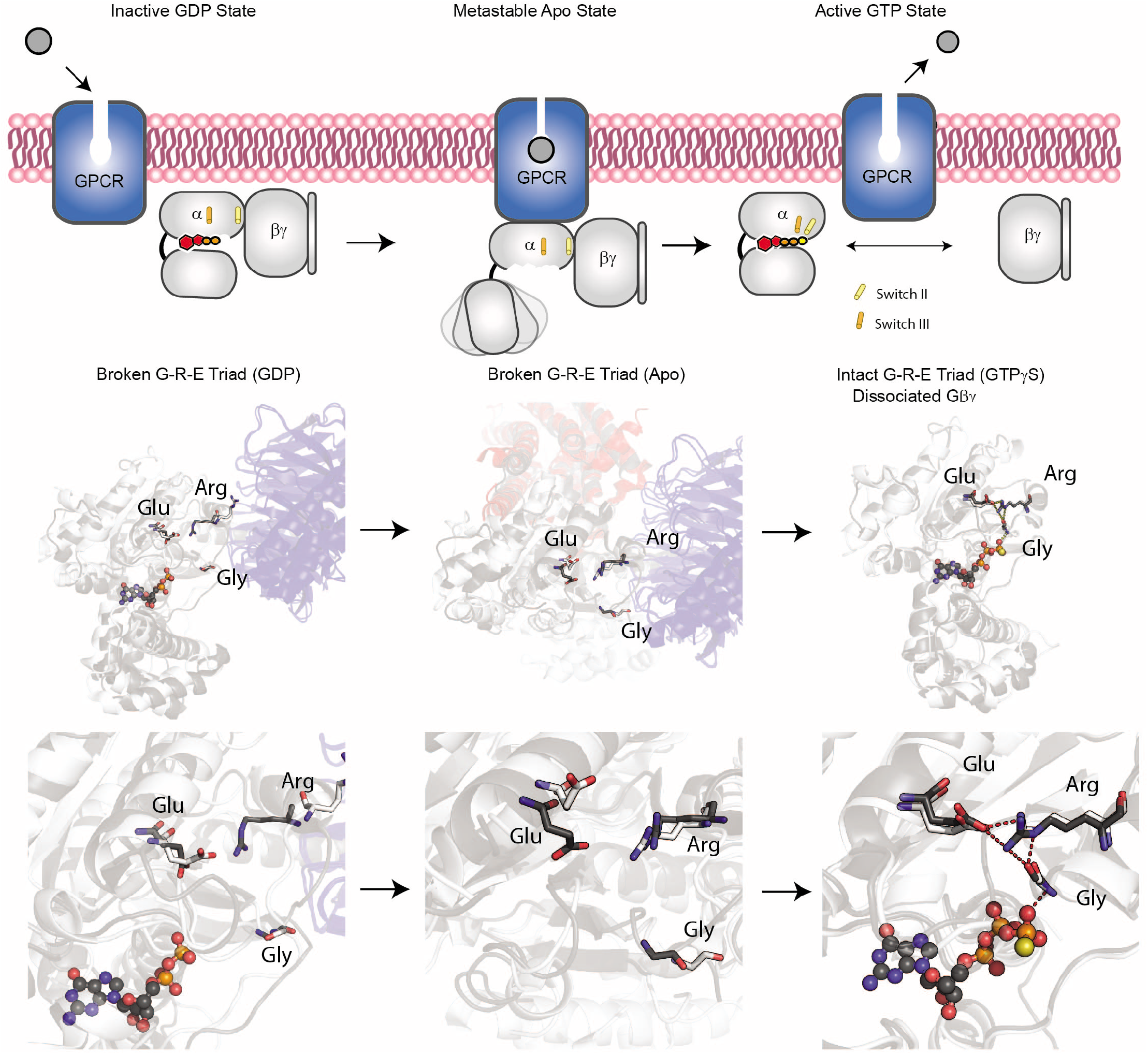
Schematic of the G protein triad hypothesis. **Top**, schematic representation of the G protein (grey) bound to GDP (left), none (center), and GTP (right). Cylinders represent the movements of Switch II and Switch III. **Bottom**, overlaid structural representation of Gα_i1_ and Gα_s_ (grey) bound to GDP (PDB: 1GP2 6CRK) (Maeda et al., 2018; Wall et al., 1995), none (apo state, PDB:6DDE 3SN6) (Koehl et al., 2018; Rasmussen et al., 2011), and GTP (PDB:1GIA and 1AZT) (Coleman et al., 1994; Sunahara et al., 1997). The overlaid residues for the conserved triad are highlighted in black.

To guide our experimental analysis, we began with a detailed examination of the known Gα structures. In particular we focused on the amino acids that bridge the γ-phosphate to the Gβγ binding domain. This analysis revealed a triad of amino acids (in Gα_i1_ numbering: G203, R208, and E245) that are conserved in all Gα proteins (Figure S1, Supplemental Information). On GTP binding, the glycine repositions the arginine and allows it to form a salt bridge with the glutamate. This Arg-Glu pair has been described previously as a molecular “hasp” or latch that fastens Switch II to the α3 helix and locks Gα in the active conformation (Figure 1) (Iiri et al., 1997). Accordingly, the conserved glycine functions as a lock or pin that secures the latch and prevents access of Gβγ to Gα. In support of this model, small G proteins such as RAS contain the conserved glycine but lack the arginine or glutamate and do not bind to Gβγ subunits. Moreover, replacement of the conserved glycine in Gα_s_ (G226A) or Gα_i1_ (G203A) disrupts formation of the Glu-Arg salt bridge (Berghuis et al., 1996), and appears to stabilize the protein in the Gβγ-bound state (Berghuis et al., 1996; Lee et al., 1992; Miller et al., 1988). Mutations in the arginine (Gα_s_-R231H and yeast Gpa1-R327S) likewise impede cell signaling by Gβγ (Apanovitch et al., 1998; Iiri et al., 1997). Based on our modeling and previous observations, we hypothesized that this triad of amino acids senses the presence of the γ-phosphate of GTP and allosterically controls subunit dissociation.

### Triad residues regulate Gβγ signaling in vivo

While RAS and Gα are functionally related and share the same architecture, the acquisition of the arginine-glutamate latch may account for the ability of Gα to regulate Gβγ release. As an initial test of our model, we replaced the conserved arginine and glutamate with all 19 other amino acids and tested their functionality in a systematic fashion. Given the large number of mutants and, given the need to isolate the effects of each mutant from other functionally similar Gα subtypes, we conducted our analysis in yeast. The yeast system has some unique advantages that would help in our interpretation of the data. Most notably, yeast express a single canonical GPCR and G protein, greatly simplifying analysis of structure-function relationships. This is in marked contrast to humans where multiple G proteins receive signals from hundreds of distinct receptors. Moreover, G proteins in yeast and animals have conserved structures, and heterologous expression studies indicate that the receptors and G protein subunits are functionally interchangeable (Dowell and Brown, 2009). Therefore, any perturbation in the yeast Gα are usually predictive of perturbations in mammalian G proteins.

We first compared pheromone signaling in cells that express wild-type or mutant forms of Gpa1, expressed using the native promoter in a single copy plasmid. All of our functional assays used the extensively-studied G323A mutant as a positive control. This substitution has been characterized mechanistically for Gα_s_ and Gα_i1_, and demonstrated to reduce the binding affinity for magnesium and GTP (Berghuis et al., 1996; Lee et al., 1992; Miller et al., 1988). These mutants bind properly to receptors, GDP-Pi and Gβγ, and partially stimulate (25-60% of full activity) adenylyl cyclase in vitro. Thus, the Gly-to-Ala mutants appear to be locked in a permanently Gβγ-bound state, but are otherwise properly folded and competent to interact with other known binding partners. Based on these previous studies, and given the proximity of Gly323 to the guanine nucleotide, we predicted that G323A would exhibit the strongest suppression of G protein dissociation, making it our benchmark control.

We measured two outcomes of pheromone signaling. The first was Gβγ-mediated cell growth arrest (halo assay). Pheromone spotted onto a filter disk produces a zone of growth arrest, the size of which correlates with pheromone sensitivity. As shown in Figure 2, substitutions of Arg327 with Cys, Asn, Ser, and Val conferred the most effective inhibition (Figure 2A, bottom). Inhibition was absent when the arginine was replaced with glutamine. Other replacements yielded intermediate results. Most substitutions of the triad glutamate likewise yielded intermediate effects, with the strongest effect observed for E364K. None of the arginine or glutamate substitutions were as potent as the benchmark G323A mutant.

**Figure 2.**
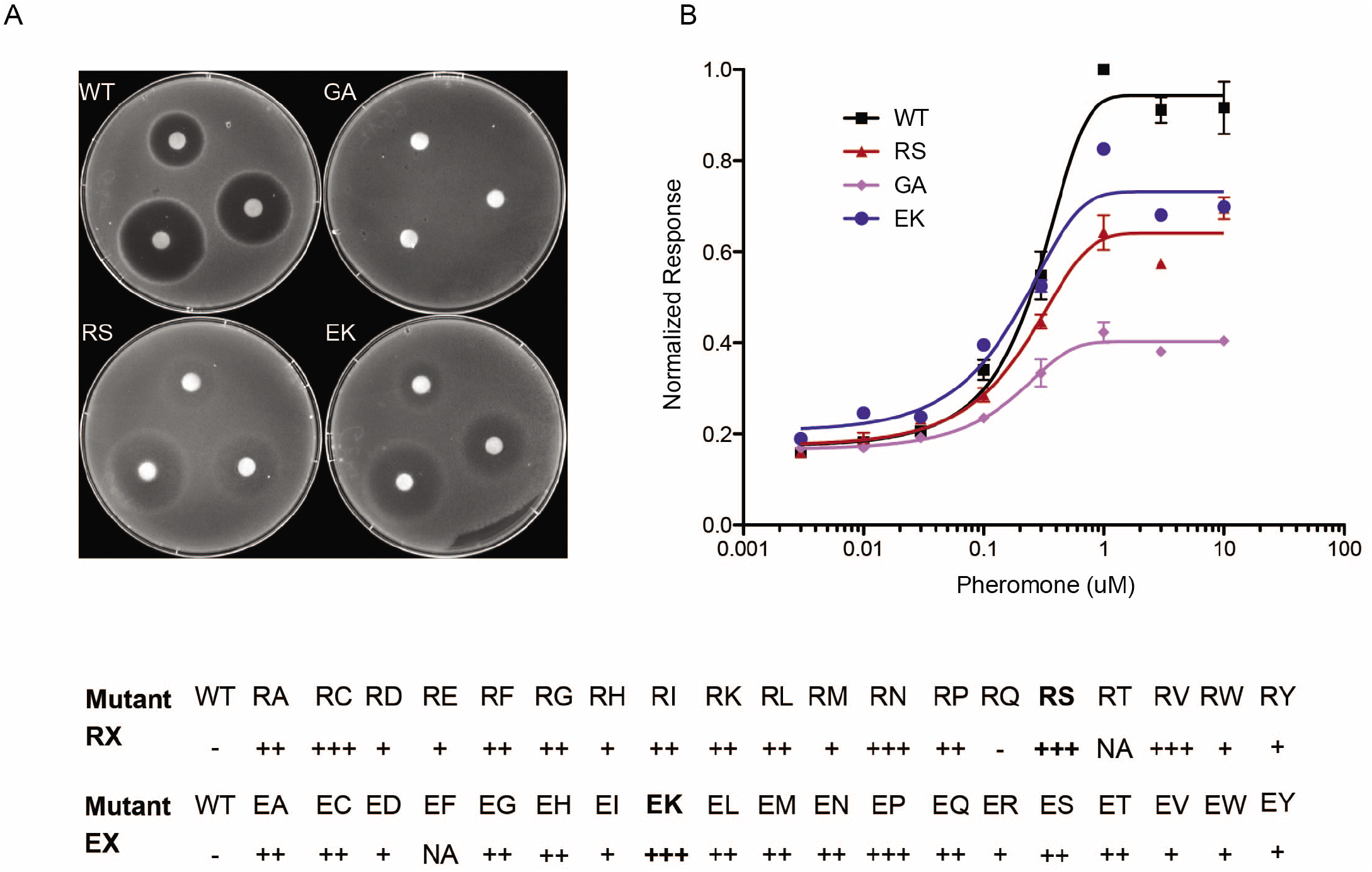
Triad residues regulate Gβγ signaling in vivo. Wild-type yeast cells transformed with a single copy plasmid containing *GPA1* or the indicated mutant G323A (GA), R327S (RS) or E364K (EK). **Left**, plate assay with filter disks containing 75, 25, or 8 μg α-factor pheromone. **Right**, transcription reporter assay after treatment with the indicated concentration of α factor pheromone. **Bottom**, quantitation of mutant phenotypes. Values indicate extent of growth inhibition, ranging from least (-) to most (+++) turbidity.

We then examined the effects of the triad mutations using a Gβγ-dependent gene transcription assay. In this method, induction of a pheromone-inducible promoter (from *FUS1*) leads to increased expression of a reporter protein. As shown in Figure 2B, expression of Gpa1-R327S caused a substantial reduction in reporter activity, but not to the extent of the benchmark G323A mutant. Thus, the two functional assays are largely in agreement and together show that pheromone signaling is diminished upon expression of some, but not all, triad mutants. Given that almost any substitution should disrupt the salt bridge that defines the hasp, we conclude that arginine and glutamate have unique functional properties that go beyond their ion-pair interaction.

### Triad residues control subunit dissociation

Our yeast-based measurements indicate that many, but not all, triad substitutions exhibit inhibitory effects on Gβγ release. To determine if these functions are universally conserved, we introduced our best-performing mutations into human Gα subunits and tested their activity in a mammalian expression system. To this end we used an optimized protein proximity assay that detects receptor-stimulated release of Gβγ from Gα. In this bioluminescence resonance energy transfer (BRET) assay, a Gα-*Renilla* luciferase (RLuc) fusion is co-transfected with Gβ_3_ and a Gγ_9_-GFP fusion (Olsen et al., 2020). For these experiments we used the neurotensin receptor NTR1 because it couples efficiently to multiple Gα subtypes including Gα_i_, Gα_q_ and Gα_s_. When agonist stimulates the receptor, dissociation of the Gα-RLuc (energy donor) and Gβγ-GFP (energy acceptor) is measured as the ratio of the emission of GFP to the emission of RLuc. As shown in Figure 3A, activation of all three G proteins was substantially diminished by the triad substitutions, with the exception of the Arg-to-Gln replacement in Gα_i3_, (but not Gα_s_ or Gα_q_, discussed below). Signaling by the μ-opioid receptor was likewise dampened by mutations in Gα_i3_, indicating that the inhibitory effects are shared by multiple receptors and agonists (Figure S3, Supplemental Information). We conclude from these data that the triad residues are not only structurally conserved but are functionally conserved as well. These findings also demonstrate the utility of the yeast system for prioritizing mutants suitable for follow up analysis in a more complex but physiologically informative system.

**Figure 3.**
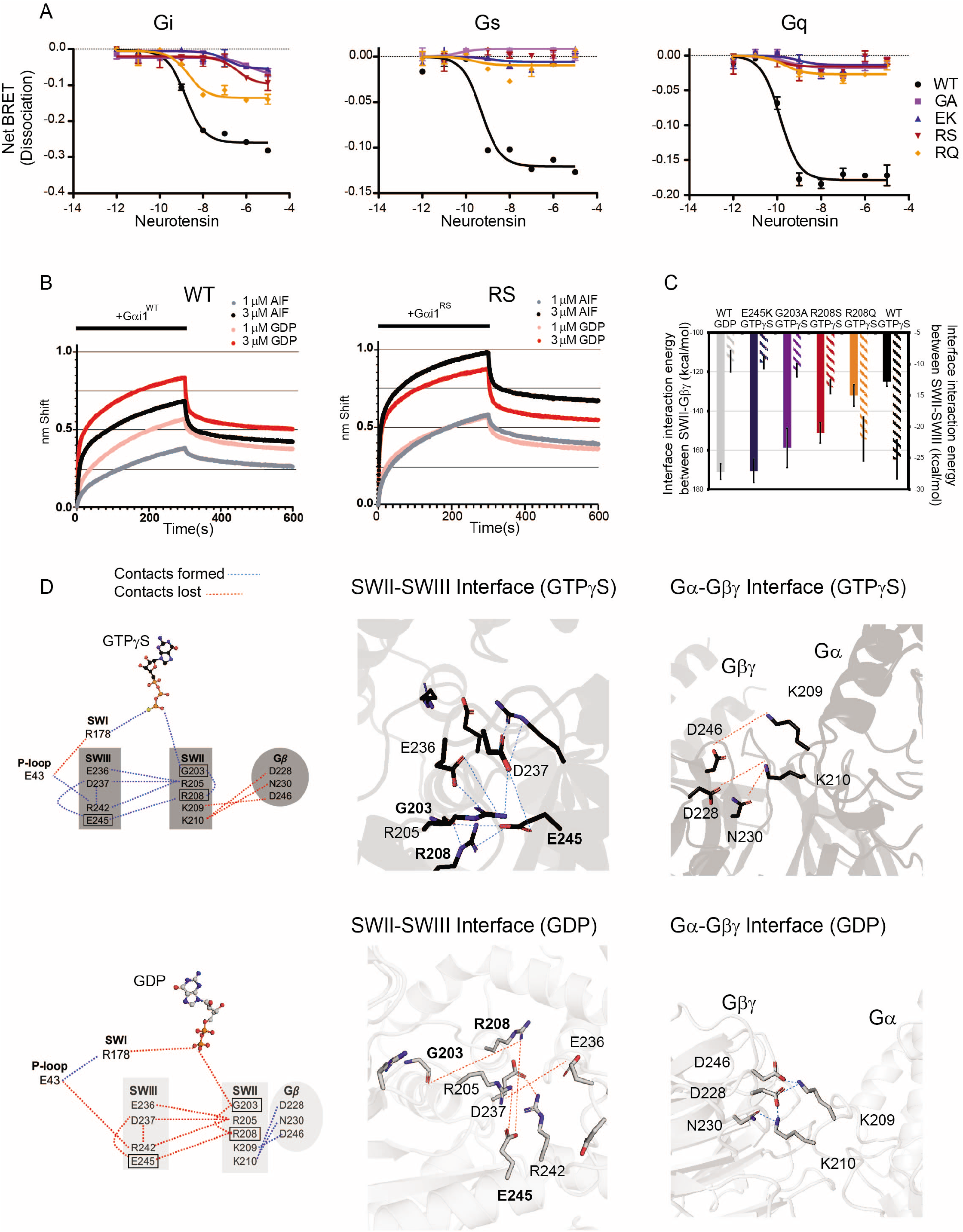
Triad residues control subunit dissociation. **(A)** HEK293 cells transfected with the indicated wildtype or mutant Gα-RLuc8 donor and Gγ-GFP acceptor proteins. Concentration-response measurements (n=2) using the neurotensin receptor are presented as fold-decrease in dynamic range (dissociation, Net BRET). **(B)** Purified biotinylated-Gβ and Gγ immobilized on streptavidin exposed for 300 sec (bar) to the indicated concentration of purified Gα_i1_ (WT, left) or Gα_i1_-R208S (RS, right), equilibrated in either GDP (red or pink) or GDP and aluminum fluoride (AlF_4_^-^, black or grey). Binding is reported as a shift in the interference pattern (nm). **(C)** Plot of interaction energies between Gα and Gβ subunits versus interaction energies between residues in Switch II and Switch III regions in wildtype and the indicated Gα_i1_ mutants. The interaction energies are averaged over the snapshots of the MD simulation trajectories. **(D)** A dynamic mechanism for the effect of the γ-phosphate group triggering the release of the Gβγ subunits. The dynamic pull-push effect from the γ-phosphate group of GTPγS, that forms the polar residue interaction network involving the P-loop, Switch I, Switch II and Switch III residues, derived from MD simulations for wildtype Gα_i1_ bound to GTPγS (top) or GDP (bottom). The nucleotides are shown in ball and stick representation. The blue and the red dotted lines indicate sustained and broken residue contacts, respectively. Right panels show residues involved in the polar network for the GTPγS- and GDP-bound Gα_i1_ switch regions (black and gray colors respectively).

The data provided above indicate that the conserved triad is needed for proper Gβγ release in yeast and in human cells. To delineate the mechanism by which the triad acts, we tested the ability of the R208S mutant form of the protein to bind Gβγ directly. We focused on the conserved arginine because it connects the Switch II glycine (binds γ-phosphate) and the Gβγ binding interface. In addition, the R208S mutant exhibited the strongest inhibitory effect of the 19 substitutions tested. For this analysis we chose bio-layer interferometry (BLI), which analyzes the interference pattern of white light reflected from two surfaces: a layer of immobilized protein on the biosensor tip, and an internal reference layer. We combined biotin-Gβ_1γ2_, immobilized onto Streptavidin-coated biosensor tips, with a solution containing either 1 μM or 3 μM purified Gα_i1_ or Gα_i1_-R208S, and equilibrated the protein in either GDP or GDP plus aluminum fluoride (AlF_4_^-^). Aluminum fluoride mimics the pentavalent transition state for GTP hydrolysis (Mixon et al., 1995; Sondek et al., 1994). Thus, the structure of the AlF_4_^-^ state is very similar to the active GTP-bound state and preserves the triad polar network, but activation by AlF_4_^-^ does not depend on nucleotide exchange. After 5 min the tips were transferred to an identical solution but lacking Gα_i1_. By this measure, the wildtype (Figure 3B, left) and R208S mutant (Figure 3B, right) exhibited similar association and dissociation kinetics in the presence of GDP (red and pink traces) (Table S1, Supplemental Information). As expected, GDP-AlF_4_^-^- Gα_i1_ bound more poorly to Gβγ (black and grey traces), with a slower initial on rate and lower maximum binding at both concentrations of Gα subunit. In contrast, and in support of our model, Gβγ bound equally well to the “active” and “inactive” forms of the R208S mutant. The lack of an AlF_4_^-^-dependent alteration of the R208S binding to Gβγ is consistent with the BRET data obtained in HEK293 cells (Figure 3A). Thus, the conserved arginine mediates the assembly of the triad and formation of an active state conformation leading to a lower affinity of Gα for Gβγ.

### A dynamic mechanism for the γ-phosphate of GTP in coordinating the active-state conformation of Gα

To provide a dynamic view of events leading to dissociation of the Gβγ subunits, we performed all-atom MD simulations of Gα_i1_ in complex with Gβγ and bound to either GDP or GTPγS. We also simulated the dynamics of Gα_i1_-R208S, Gα_i1_-E245K and Gα_i1_-G203A, all bound to GTPγS. Finally, as a counter example to these loss-of-signaling mutants, we included in our analysis Gα_i1_-R208Q, which is unique in its ability to retain partial activity. Our analysis of the crystal structures of the GDP-bound trimeric Gα_i1_Gβγ (Wall et al., 1995) compared to GTPγS-bound Gα_i1_ (Coleman et al., 1994) showed that the residues in Switch II and Switch III form tighter interactions when bound to GTPγS and enable Gβγ dissociation. To assess how mutants affect the strength of subunit interaction, we calculated the non-bond interaction energies between the Gα and the Gβ subunits, averaged over MD trajectories for each of the mutant and wild type proteins. We also calculated the interaction energies between the residues in the Switch II and the Switch III regions (see Methods section for more details). As anticipated, the Gα-Gβ subunit interaction energies varied inversely with the interaction energy between the Switch II and Switch III regions, shown in the plot in Figure 3C. In contrast, the non-signaling Gα_i1_ mutants showed Gα-Gβ interaction energies similar to that of GDP-bound wildtype Gα_i1_ even when bound to GTPγS.

To understand the mechanism of this dynamic “pull-push” effect we analyzed the residue interaction network emerging from the γ-phosphate of GTP connecting to Switch I, and cascading further to Switch III and the Switch II region, which forms the Gβ interface. As shown in the network diagram in Figure 3D, we identified an extensive polar network linking the γ-phosphate group to Arg178 in Switch I. This pull breaks the interaction of Arg178 with Glu43 in the P-loop, which in turn strengthens the interaction of Glu43 with Arg242 in the Switch III region. This interaction between Glu43 and Switch III leads to the triad formation, comprised of Glu245-Arg208 and Gly203 in the Switch II region. Additional polar interactions between Arg242, Asp237, Glu236 in Switch III with Arg205 in Switch II also strengthen the Switch II – Switch III interactions. This Switch III - Switch II pulling effect in turn weakens the polar network of interactions of Lys210 and Lys209 in Switch II with Asp246, Asn230 and Asp228 residues in the Gβ subunit. Breaking γ-phosphate interaction with Arg178 in the GDP-bound Gα_i1_ disrupts the cascade of polar interactions between Switch I, Switch III and Switch II, thereby breaking the triad and preserving the polar interaction network between Switch II and the Gβ interface as shown in Figure 3D. A similar interaction network mechanism was also observed for Gα_s_ MD simulations (Figure S3, Supplemental Information). These results reveal a common mechanism leading to Gβγ dissociation. Specifically, we have identified an allosteric link between the terminal phosphate of GTP and the Gα-Gβγ interface.

### Triad mutants confer a GDP-like conformation on GTP-bound Gα_i1_

Our cell-based assays indicate that mutations in the triad arginine and glutamate confer sustained binding to Gβγ. The experimental BRET (Figure 3A) agrees reasonably well with the subunit interface interaction energy calculated from MD simulations. Our MD simulations suggested that these mutants lack important conformational changes associated with GTP binding, specifically changes in Switch II and Switch III, and that these structural differences are propagated by the γ-phosphate of GTP. To that end, we monitored the conformational stability of all three mutants when bound to guanine nucleotides.

We have previously shown that Gα_i1_ is thermodynamically more stable when bound to GTPγS than to GDP (Isom et al., 2013). This difference is consistent with structural studies showing that the GTP-bound state is considerably more rigid than the GDP-bound state, as dictated by regions near the γ-phosphate group and at the Gβγ interface (Sun et al., 2015). Since the triad mutants function as if they were bound to GDP, we anticipated that they would exhibit a reduced thermostability even if bound to GTP. To test this, we purified wildtype and several mutant forms of Gα_i1_ and determined the melting temperature of each using the quantitative cysteine reactivity (fQCR) assay. In this method, cysteine residues are labeled with a fluorogenic reagent by a mechanism that is analogous to hydrogen exchange. Upon heating, protected cysteine residues of Gα become exposed and can be covalently labeled by cysteine-specific probes (Isom et al., 2013). Using a two-state model of denaturation, we analyzed each Gα_i1_ mutant unfolding profile to quantify the midpoint of temperature unfolding (*T*_m_). Representative thermostability profiles for the wildtype and triad mutant forms of Gα_i1_ are shown in Figure 4A. These experiments confirm that the wildtype protein undergoes a 9 °C increase in *T*_m_ when bound to GTPγS in place of GDP; in contrast, the mutants exhibited little or no change. We conclude that that the conserved triad confers critical conformational changes that favor the dissociated state of the G protein.

**Figure 4.**
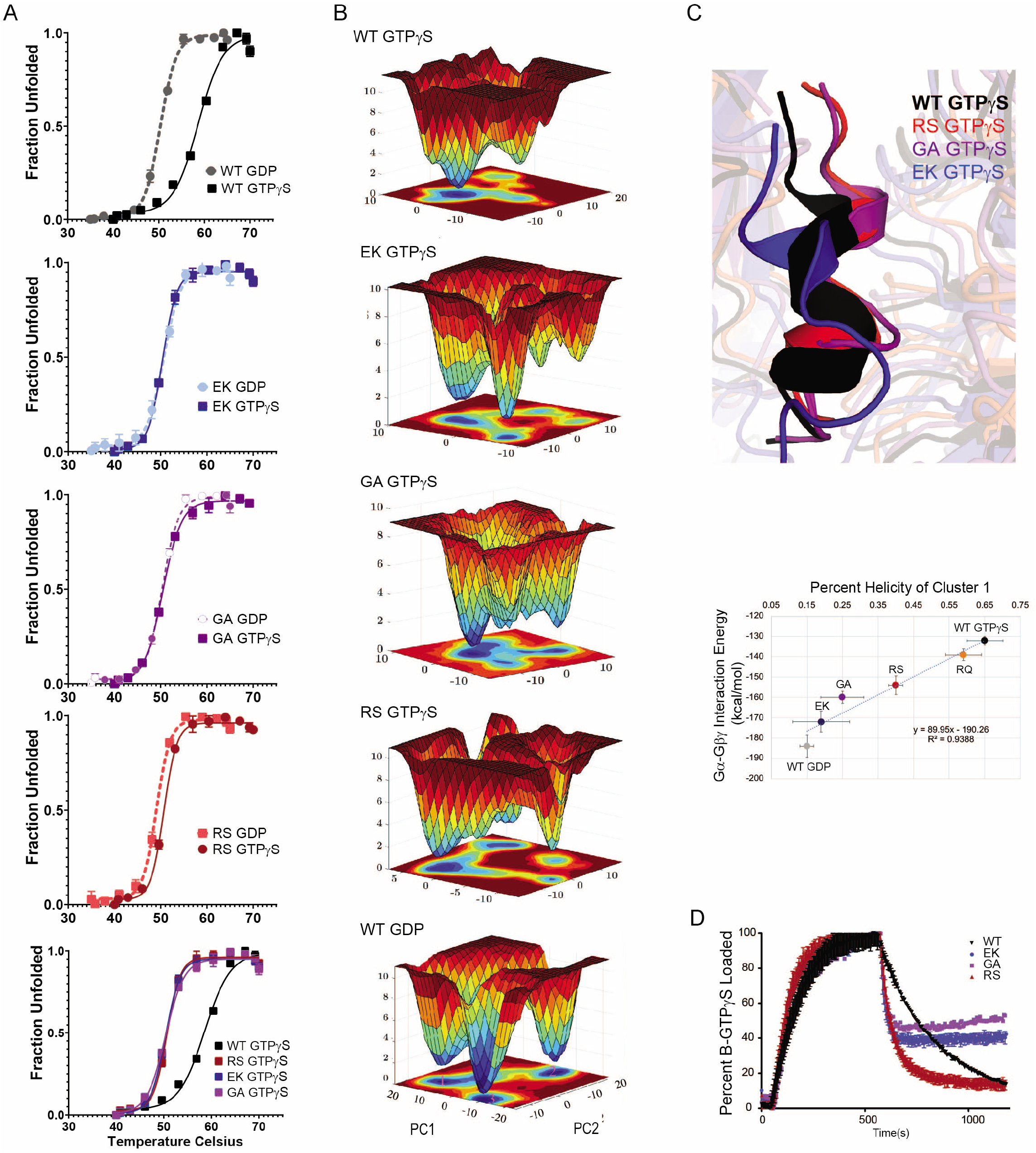
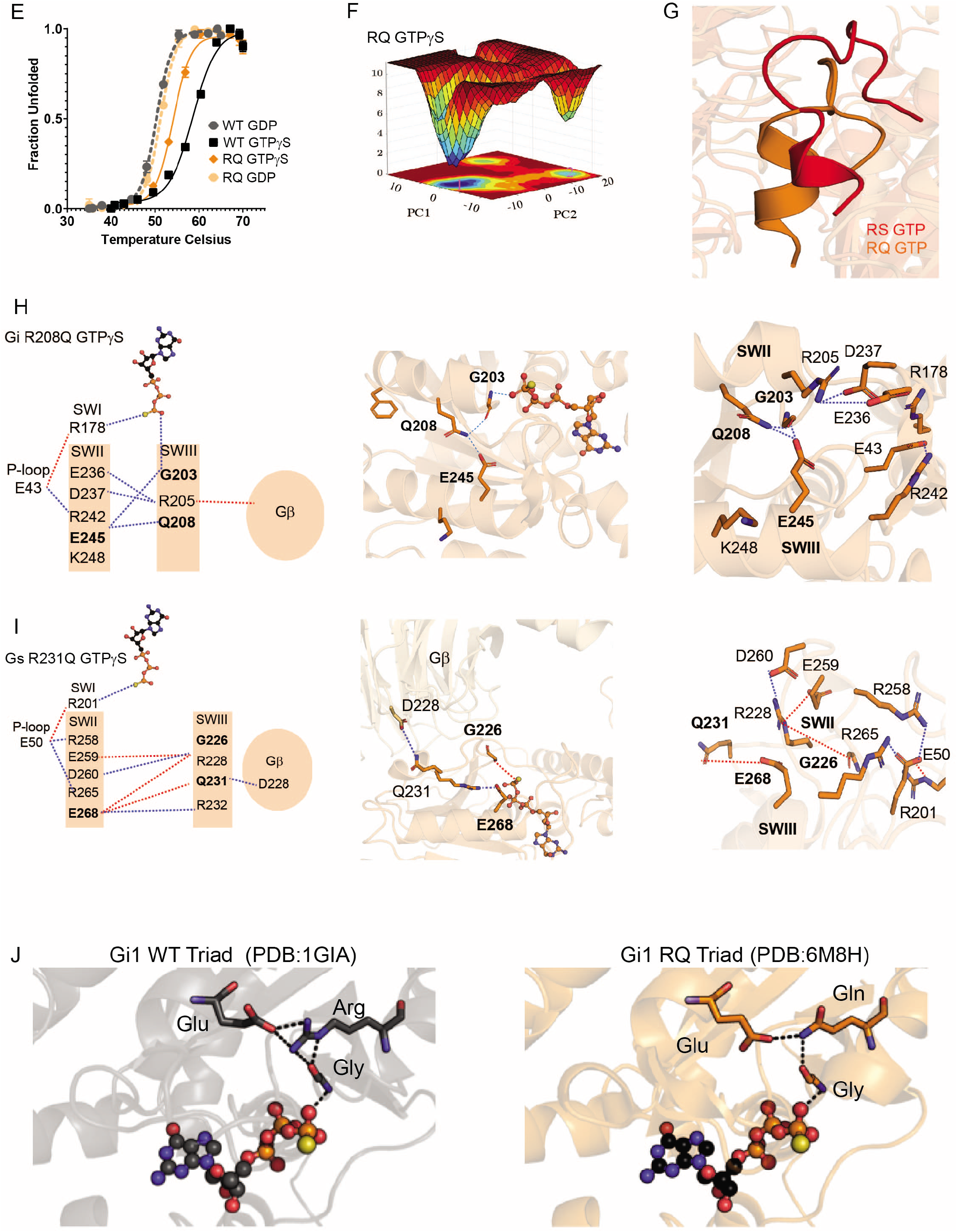
Triad residues coordinate the active-state conformation. **(A**) Thermostability of purified wildtype and the indicated Gα_i1_ mutants equilibrated in either GTPγS (solid lines) or GDP (dashed lines). *T*_m_ values were quantified by fitting a two-state model of thermal unfolding (n=3). **(B)** Free energy surfaces computed using population densities show the conformational heterogeneity of Gα_i1_ and its mutant systems. The projections of the surfaces are shown on the principal components 1 and 2 (PC1 and PC2) axes. The various conformational clusters are numbered (magenta) ordered by the population of each conformation cluster. Cluster 1 for example, is the most populated conformation cluster. The Z axis is the Gibb’s free energy obtained by inverting the population of these microstates and is shown as a colored heatmap. **(C)** Helicity of residues in the Switch II regions of wild type and the indicated Gα_i1_ mutants, bound to GTPγS, calculated as an average over the snapshots in the most occupied conformational cluster I from the free energy surfaces. **(D)** Association of 6 μM purified wildtype and the indicated Gα_i1_ mutants with 100 nM BODIPY-GTPγS, followed after 600 s by addition of 50 μM unlabeled GTPγS (n=2). **(E**) Thermostability of purified wildtype Gα_i1_ and R208S mutant, as detailed above (A). **(F)** Free energy surfaces computed using population densities calculated for principal components PC1 and PC2 for Gα_i1_-R208S, as detailed above (B). **(G)** Structural representation of the Switch II regions of Gα_i1_-R208S bound to GTPγS, as detailed above (C). **(H)** A dynamic mechanism for the effect of the γ-phosphate group triggering the release of the Gβγ subunits for Gα_i1_-R208Q as detailed above in Figure 3D. **(I)** A dynamic mechanism for the effect of the γ-phosphate group triggering the release of the Gβγ subunits for Gα_s_-R231Q as detailed above in Figure 3D. **(J)** Crystal model of Gα_i1_ and Gα_i1_-R208Q triad residues and GTPγS (ball and stick) from PDB: 1GIA and 6M8H.

Structural dynamics studies have shown that thermostable mutants exhibit conformational homogeneity, meaning fewer distinct conformations in the ensemble during dynamics simulations (Vaidehi et al., 2016). The conformational homogeneity manifests as fewer energy minima in the free energy surface of thermostable mutants (Ghosh et al., 2018). To understand the structural basis for the observed thermostability differences, we generated the free energy surface of GDP- and GTPγS-bound Gα_i1_ using MD simulation trajectories. Figure 4B shows the free energy surface of Gα_i1_ bound to GTPγS and GDP, along with the Gα_i1_ mutants bound to GTPγS. In particular, we observed that Gα_i1_-GTPγS had fewer energy minima and was less conformationally homogeneous compared to the mutant forms of Gα_i1_. Thus, there is a correlation between conformational heterogeneity and thermal instability.

A comparison of crystal structures of GDP- and GTPγS-bound Gα_i1_ shows that residues in Switch II fold into a helix leading to the dissociation of Gβγ. We speculated that formation of an ordered helical region in Switch II in GTPγS-bound Gα_i1_ could lead to its enhanced thermostability. Therefore, we calculated the average helicity of the residues in the Switch II region averaged over all the MD snapshots in the most occupied conformation cluster extracted from the free energy surfaces for the wildtype and mutant forms of Gα_i1_. As shown in Figure 4C, Switch II transitioned from about 15% to 63% helicity in going from GDP-bound to GTPγS-bound Gα_i1_. It is important to note that we started the MD simulations from the same disordered conformation in Switch II observed in the crystal structure of the GDP-bound Gα_i1_ trimeric structure. MD simulations recapitulate the loop-to-helical transitions that are triggered by the γ-phosphate group of the GTPγS engaging the dynamic mechanism shown in Figure 3D. To examine if the loop-to-helix transitions lead to increased thermostability, we calculated the average interaction energies between the Gα and Gβ subunits averaged over the MD snapshots in the most occupied conformation cluster, extracted from free energy surfaces. The calculated helicity of Switch II correlates inversely with the Gβγ interaction energies for wildtype Gα_i1_, bound to GDP and GTPγS, as well as the Gα_i1_ mutants bound to GTPγS (Figure 4C). In summary, these findings indicate that increased helicity could bring a more ordered structure to the Switch II region and confer thermostability and conformational homogeneity for Gα_i1_ bound to GTPγS.

An alternative interpretation is that the mutants bind to GDP but not GTP. To test this experimentally we determined the rate of nucleotide exchange, measured as a gain of fluorescence as BODIPY-GTPγS bound to protein (McEwen et al., 2001). As shown in Figure 4D, the labeled analog bound to mutants as well (or faster) than the wildtype protein. We then added an excess of unlabeled GTPγS and determined that the rate of dissociation was substantially faster for all of the mutants as compared to that of wildtype. We infer that the triad mutants confer a more dynamic binding pocket, which results in faster dissociation of nucleotide. More broadly, these data indicate that loss of the triad residues does not impede GTP binding or dissociation, but prevents the conformational changes required for Gα activation and Gβγ dissociation. Taken together, our MD simulations and biophysical assays confirm that mutations in the triad arginine and glutamate place the protein in a permanently “inactive” conformation, compatible with sustained binding to Gβγ.

The results presented above reveal the unique importance of the triad arginine. Generally speaking, substitutions at this site lead to diminished signaling by Gβγ. However, as detailed above (Figures 3 and 4), substitution of the arginine with glutamine partially preserves function, but only in Gα_i1_ and not in Gα_s_. In accordance with these observations, our calculations of interaction energies (Figure 4C), thermal stability measurements (Figure 4E), computed free energy surfaces (Figure 4F), and helicity calculations (Figures 4C and G) revealed that the R208Q mutant is more similar to wildtype than any of the other arginine mutants. The weakening of interaction between Gα_i1_ and the Gβγ subunits of G protein is caused by the enhanced interaction between the Switch II and Switch III regions within the Gα_i1_ subunit. These simulations are the theoretical basis for the proposed competition between Gβγ and Switch III for Switch II (Figure 4H). The calculated interaction energies for Gα and Gβγ subunits become less favorable as the interaction energies for Switch II and Switch III become more favorable, and vice versa (primary and secondary y-axis respectively, Figure 3C). The same paradigm holds for Gα_s_ (Figure 4I). Thus, our molecular dynamics simulations were able to predict substitution- and subtype-specific dissociation events with exquisite accuracy and mechanistic detail.

We then sought to determine the structural basis for the unique ability of Gα_i1_-R208Q to sustain signaling. To that end, we compared the structure of wild type Gα_i1_ with a new X-ray crystal structure of Gα_i1_-R208Q, both in the GTPγS-bound state (Figure 4J, compare left and right panels). This analysis revealed that Gln208, like the native arginine of Gα_i1_, is close enough to form a H-bond network with the conserved glycine and glutamate. In contrast, and as anticipated by our MD simulations, the same substitution would not restore the network in Gα_s_ or Gα_q_. Whereas Arg208 of Gα_i1_ faces Switch III in both the active and inactive states, the corresponding residue in Gα_s_ must rotate ~180° before it can interact with the conserved glycine and glutamate.

Our simulations appear to account for the distinct functional properties of the G-R-E motif, as determined by thermostability measurements, nucleotide and protein binding measurements, as well as the signaling outputs obtained in yeast and animal cells. Most strikingly, the atomistic MD simulations anticipated the unique and unexpected differences between G_i_ and G_s_, and the differences between R208Q and other Arg208 substitutions. The differences between Gα_i1_-R208Q and Gα_s_-R231Q in particular support our model that the arginine has functions beyond simply forming a salt bridge with the conserved glutamate. More broadly, these results highlight the predictive power of MD simulations, particularly when integrated with molecular and cellular experimental analysis.

### The triad arginine controls the final committed step of G protein activation

Finally, we sought to determine the structural basis for the unique functional properties of the R208S mutation. Based on the mechanism from MD simulations, we postulated that the mutant retains Gβγ binding by disrupting the conserved polar network that dictates the response to GTP. To test this directly, we collected ^1^H-^15^N 2D heteronuclear NMR spectra of wild-type and mutant Gα_i1_, both in the presence of GDP and GTPγS. This method allows for the detection of backbone and side-chain NH resonances. As an NH resonance can be detected for every residue with the exception of proline, the spectrum contains a “fingerprint” of the protein backbone and perturbations resulting from changes in intramolecular interactions. We consider this a definitive method for detecting the conformational changes that accompany nucleotide binding and G protein activation. In contrast to some other Gα_i_soforms and most members of the RAS family of GTPases, Gα_i1_ undergoes nucleotide exchange within minutes, even in the absence of exchange factor or receptor (Figure 4D).

As shown previously (Goricanec et al., 2016), a substantial number of Switch II peaks (Figure 5 arrows) for wildtype Gα_i1_ were resolved in the presence of GTPγS (black) but not GDP (blue). These data are consistent with X-ray structural data, indicating that Switch II is more ordered in the activated state as compared with the inactive state. Likewise, Gα_i1_-R208S exhibited multiple peak shifts when GDP was replaced with GTPγS, indicating that nucleotide exchange had occurred (Figure S5 top, Supplemental Information). Consistent with our fast exchange data for R208S, the mutant GTPγS spectrum contains a shifted tryptophan that indicates GTPγS binding (Figure 5A circle). In contrast to wildtype, Gα_i1_-R208S appears to be missing multiple peaks, consistent with a more dynamic GTPγS state. In support of our observations from MD simulations (Figure S3 Supplemental Information), many resonances missing in the mutant spectrum were in the Switch regions for wildtype bound to GTPγS (Figure 5B). These results indicate that Gα_i1_-R208S is in an alternate, more dynamic conformational ensemble, even when bound to GTPγS. More broadly, these results indicate that the conserved arginine is necessary to detect the presence of the γ-phosphate and undergo the structural changes necessary to maintain the active state.

**Figure 5.**
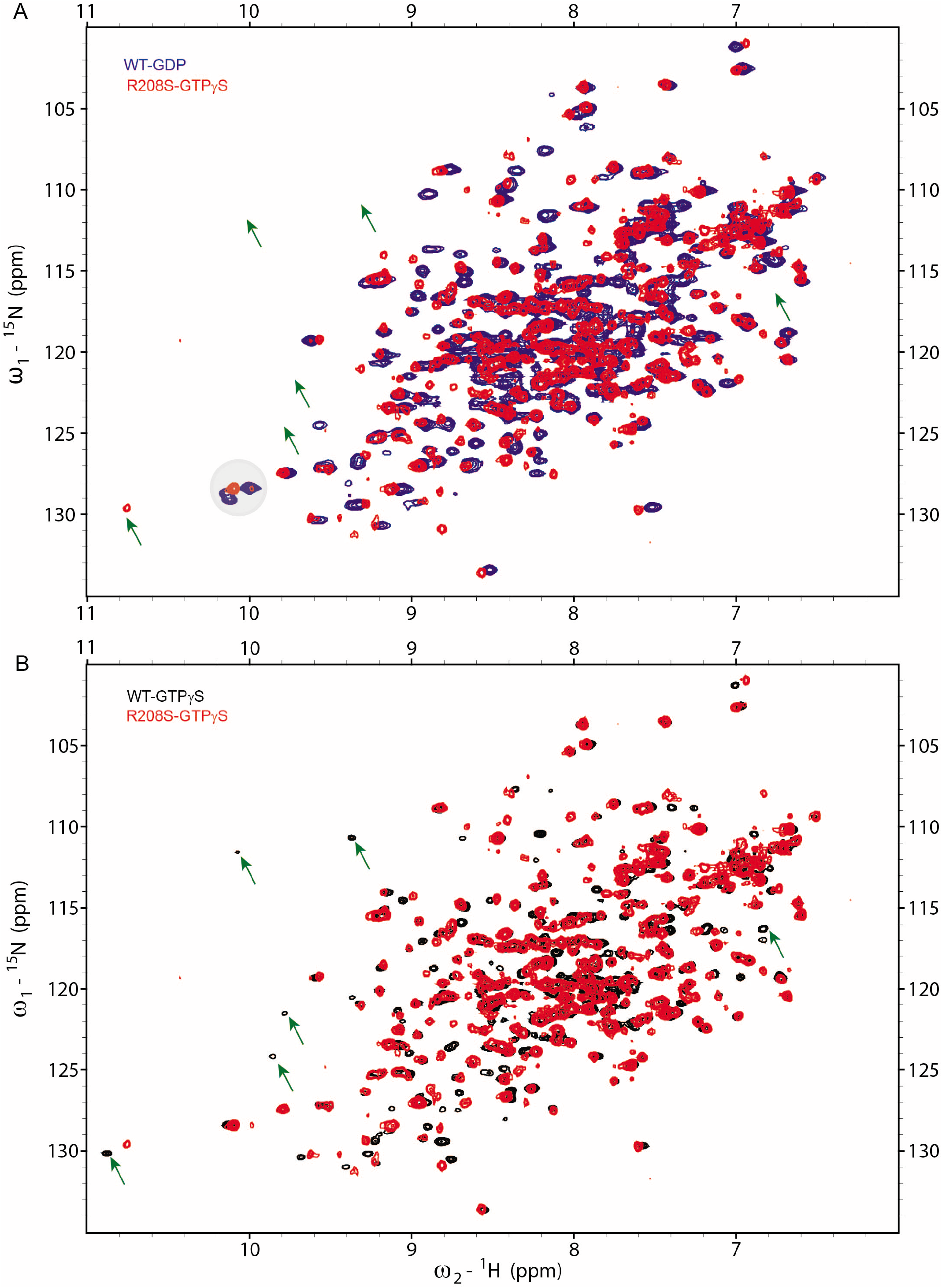
The triad arginine controls the final committed step of G protein activation. 2D [^15^N, ^1^H]-HSQC NMR of Gα_i1_Δ31 wildtype and R208S mutant. **(A)** Overlay of wildtype Gα_i1_Δ31 in complex with GDP (Blue) and R208S Gα_i1_ in complex with GTPγS (Red). **(B)** Overlay of wildtype (Black) and R208S (Red) Gα_i1_, each in complex with GTPγS (Black). Switch II specific peaks are labeled (arrows).

## DISCUSSION

Heterotrimeric G proteins are present in a wide variety of organisms and exist in multiple distinct subclasses. Recent findings, including structure determinations of receptor-bound G proteins, have revealed the molecular motions leading to the release of GDP. Here we considered the subsequent molecular events leading to the release of Gβγ. Our approach combined crossspecies sequence analysis, cell-based functional assays, MD simulations, thermostability measurements, direct Gα and Gβγ binding measurements, and NMR spectroscopy.

Our study revealed the existence of a network of highly conserved amino acids (“G-R-E motif”) that dictates the release of Gβγ by Gα. In the presence of the terminal phosphate of GTP, the conserved glycine forms a polar network with the arginine and glutamate, disrupting the Gα-Gβγ binding interface. By systematically replacing triad residues, and measuring the functional consequences of those changes, we demonstrated its unique importance in G protein activation in cells. Whereas a single phosphate can initiate subunit dissociation, we show that mutation of a single amino acid can override the process entirely. We then determined the molecular basis of activity through biochemical and biophysical experiments conducted in vitro. Finally, we established the atomic interactions responsible for dissociation of the G protein subunits in silico.

Our findings provide the molecular basis for important functional behaviors documented through cellular genetic studies conducted almost 40 years ago. The S49 cyc^-^ cell line, which bears the Gα_s_-G226A mutation and is deficient in cAMP production (Bourne et al., 1981; Salomon and Bourne, 1981), was instrumental in the discovery of G proteins (Gilman, 1987). In yeast, spontaneous mutations at the triad glycine, glutamate and arginine confer strong resistance to pheromone stimulation (Apanovitch et al., 1998; Stratton et al., 1996). Replacement of the arginine was even sufficient to block unrestrained signaling in the complete absence of GTP hydrolysis (Apanovitch et al., 1998). The importance of the triad arginine has also been revealed through human genetic studies. The conserved arginine is one of two sites in Gα_s_ mutated in patients with pseudohypoparathyroidism 1a, a syndrome characterized by Albright hereditary osteodystrophy and resistance to hormone stimulation (Farfel et al., 1996; Iiri et al., 1997). More recent studies indicate that the triad arginine and glutamate are mutation “hotspots” in Gα_o_, frequently altered in individuals with neurodevelopmental disorders including involuntary movement and seizures. Of more than 30 missense or codon deletion mutations identified so far, at least six affect the conserved arginine or glutamate (https://monarchinitiative.org/gene/HGNC:4389#overview). All of the phenotypes associated with loss of the G-R-E motif, in yeast and humans, can be attributed to sustained subunit association.

Finally, our analysis of G protein activation complements recent advances in our understanding of receptor activation. The first crystal structure of a receptor-G protein complex, published in 2011 (Rasmussen et al., 2011), provided detailed insights into the conformational changes leading to nucleotide release. Most notably, that structure revealed a dramatic displacement of the Gα helical domain from the RAS-like domain, thereby exposing the nucleotide binding site. A major impediment to obtaining such structures is the difficulty of assembling an agonist-bound receptor with a nucleotide-free G protein. The first structures used fusions and single chain antibodies to stabilize this otherwise ephemeral state. Subsequent studies used variants of Gα containing a panel of up to 8 mutations, including substitutions of the triad glycine and glutamate (Draper-Joyce et al., 2018; Liang et al., 2018a; Liang et al., 2018b). Based on our findings, substitutions of the G-R-E motif are likely to be particularly useful in obtaining stabilized heterotrimer complexes suitable for structural analysis in the future.

Taken together, our analysis reveals events leading to the final committed step of G protein activation and provides a mechanistic basis for human disease. The structural rearrangements leading to subunit dissociation are more subtle than those affecting nucleotide exchange, but they are just as consequential. In both cases, conserved residues form a network of non-covalent contacts that link a bound ligand – either agonist or GTP - to a network of contacts needed to release Gβγ. Given that the G-R-E triad residues are conserved in all G proteins, the allosteric mechanism detailed here is likely to be universal for all subtypes and species. Just as a detailed understanding of the molecular basis for ligand binding has led to important advances in pharmacology, a better understanding of subunit dissociation could reveal strategies to bypass or compensate for the human genetic defects that cause disease.

## ACKNOWLEDGEMENTS

Funded by NIH grants F31NS093917 (R.H.J.O.), R35-GM134962 (S.L.C.), R35-GM127303 (A.V.S.), R01-GM117923 (N.V.), R35-GM118105 (H.G.D).

## AUTHOR CONTRIBUTIONS

Authors S.L.C., T.L., R.H.J.O., A.V.S., N.H.V., and G.Y. are listed alphabetically

K.M.K., S.G., N.V., H.G.D.: Conceptualization

K.M.K., S.G., S.L.C., T.L., R.H.J.O., A.V.S., N.H.V., G.Y., N.V., H.G.D.: Formal Analysis

S.L.C., A.V.S., N.V., and H.G.D.: Funding acquisition

K.M.K., S.G., T.L., N.H.V., G.Y.: Investigation

S.L.C., A.V.S., N.V., H.G.D.: Project administration

S.L.C., R.H.J.O., A.V.S., N.V., H.G.D.: Supervision

K.M.K., S.G., S.L.C., T.L., R.H.J.O., A.V.S., N.H.V., G.Y., N.V., and H.G.D.: Validation

K.M.K., S.G., N.V., H.G.D.: Writing – original draft

K.M.K., S.G., S.L.C., T.L., R.H.J.O., A.V.S., N.H.V., G.Y., N.V., H.G.D.: Writing – review & editing

## STAR METHODS

### Resource Availability

Lead Contact: H.G.D.

Materials Availability: The study did not generate new unique reagents

Data and Code Availability: Data are available from H.G.D. and N.V. The study did not generate new unique code. NMR data will be made available at: http://www.bmrb.wisc.edu

### Mutagenesis

Primestar Max (Takara Bio) mutagenesis was performed using primers containing 5 bp 5’ to the codon and 20-30 bp 3’ to the codon (Table S2, Supplemental Information).

### Halos

Pheromone-induced growth inhibition was measured as described previously (Hoffman et al., 2002). Briefly, 75, 25, or 8 μg synthetic α factor was spotted onto paper disks. Agar (0.5%) was melted and then kept at 55 °C before mixing with 100 μL saturated culture of *S. cerevisiae* strain BY4741 *bar1::KanMX* (*bar1*Δ), transformed with pRS316 containing either no insert (vector), *GPA1* or mutant *GPA1* (Song et al., 1996), and grown at 30 °C in selective medium. After spreading the agar/yeast mixture, each disk was applied to one of three predefined locations on the surface of the agar. Cell growth was imaged after 24 and 48 hr.

### Transcription

Pheromone-induced gene induction was monitored as described previously (Shellhammer et al., 2019). Briefly, BY4741 *bar1*Δ cells cotransformed with plasmids pRS426-pFUS1-YeGFP3 (Shellhammer et al., 2019) and pRS316-GPA1 (wildtype or mutant) were grown to saturation then diluted to OD_600_ < 0.001. After reaching OD_600_ of 0.8, synthetic α factor was added at defined concentrations. After 1.5 hr, GFP fluorescence was measured using a Molecular Devices Spectramax i3x plate reader at an excitation wavelength of 483 nm and emission wavelength of 518 nm.

### Bioluminescence Resonance Energy Transfer (BRET) assay

HEK293T cells (ATCC, #CRL-11268) were used to measure basal association and receptor-mediated dissociation of Gα and Gβγ, as described previously (Olsen et al., 2020). Briefly, cells were transfected with plasmids encoding the neurotensin or μ-opioid receptor, Gα fused to *Renilla* luciferase 8 (Rluc8), Gβ_3_ and γ_9_-fused to GFP in a 1:1:1:1 ratio (cell density 750,000 cells in 3 mL). After transfection (approximately 16 hr) cells were plated into poly-D-lysine coated 96-well white clear bottom plates in medium containing Dulbecco’s Modified Eagle Medium +1 % dialyzed fetal bovine serum and then incubated overnight. The following day, cells were washed twice with assay buffer (20 mM (4-(2-hydroxyethyl)-1-piperazineethanesulfonic acid) (HEPES), Hank’s Balanced Salt Solution, pH 7.4). 10 μL of Rluc8 substrate (coelentrazine 400a, Nanolight) was added per well at a final concentration of 5 μM. Cells were then incubated 5 min in the dark. 30 μL (3X) of agonist in drug buffer (contains assay buffer, 0.1% bovine serum albumin) was added per well then incubated for 5 min in the dark. Plates were read for luminescence at 485 nm and fluorescence emission at 530 nm for 1 sec per well using a Mithras LB940 multimode microplate reader. The BRET ratio was determined by dividing fluorescence by luminescence (GFP/Rluc). The receptor-catalyzed dissociation of the heterotrimer (net BRET) was measured by comparing the energy transfer from donor to acceptor and reported as ratios: GFP/Rluc per well - Basal BRET (GFP/Rluc at lowest dose of agonist) = Net BRET/ dissociation. The net BRET was plotted as the GFP/Rluc ratio as a function of neurotensin and the curve was fit in Graphpad Prism 8 (Graphpad Software Inc., San Diego, CA) (Che et al., 2020).

### Expression and Purification of Gα_i1_

RIPL (Agilent) or Rosetta (Novagen) cells were transformed with PET-SUMO-Gα_i1_ plasmid (Maly and Crowhurst, 2012) and grown to saturation in YZ medium containing 0.02% (w/v) glucose and 0.2% (w/v) ala-lactose; autoinduction occurred via inhibition of the Lac repressor when ala-lactose becomes the primary fuel source (Isom et al., 2013). The culture temperature was lowered from 37 to 18 °C and allowed to rotate overnight. The cultures were harvested by centrifugation for 1 hr at 4 °C and then lysed by sonication on ice. Lysate was then clarified by centrifugation for 1 hr at 4 °C. Clarified lysate was mixed with 1 mL of ProBond Ni-chelating resin (Thermo Fisher Scientific, Invitrogen #R80101) per 15 mL of lysate for 1 hr. The resin was washed twice with phosphate buffer pH 7 containing 10 μM imidazole, then SUMO-tagged Gα_i1_ was eluted using the same buffer containing 400 mM imidazole. To remove the SUMO, eluate was dialyzed at 4 °C in 4 L of phosphate buffer without imidazole after adding 1 mg ULP1 protease to the dialysis cassette. After dialysis, the cleaved product had no tag and was efficiently purified by reverse metal affinity chromatography. The protein was further purified by passing over a Sepharose Q anion exchange column (GE Healthcare) equilibrated in 100 mM potassium phosphate. The purified yield was typically 3-30 mg of Gα/L of cell culture.

### Expression and Purification of Gβ_1γ2_

High 5 cells (Invitrogen; 2 x 10^6^ cells/mL) were infected with high titer Gβ_1_ and Gγ_2_ baculoviruses. Gβ_1γ2_ was purified according to (Kozasa and Gilman, 1995), with modifications. All steps were carried out at 4 °C. Cells were harvested 60 hr postinfection by centrifugation at 2600xg and then resuspended in 50 mL of lysis buffer (20 mM HEPES, pH 8, 150 mM NaCl, 5 mM 2-mercaptoethanol (2-ME), 1 mM EDTA, 1 mL of protease inhibitor cocktail (P-2714, Sigma-Aldrich) per liter of cell culture. Cells were lysed by sonication and centrifuged at 2600xg to collect the membranes. Resuspension of membranes were accomplished by dounce homogenization in 100 mL of lysis buffer. The membranes were solubilized by adding 1% Lubrol (C12E10; Sigma-Aldrich) with stirring, and the resultant solution was clarified by ultracentrifugation at 125000xg. The supernatant was loaded onto Ni-NTA agarose (Qiagen) equilibrated with lysis buffer + 1% Lubrol. The resin was washed and the Lubrol exchanged for sodium cholate using buffers Ni-A (20 mM HEPES, pH 8, 0.4 M NaCl, 5 mM 2-ME, 0.5% Lubrol, 0.15% cholate) and Ni-B (20 mM HEPES, pH 8, 0.1 M NaCl, 5 mM 2-ME, 0.25% Lubrol, 0.3% cholate). Gβ_1γ2_ was eluted in Ni-C (20 mM HEPES, pH 8, 0.01 M NaCl, 5 mM 2-ME, 1% cholate, 200 mM imidazole). The purified yield was typically 1 mg of Gβ_1γ2_/L of cell culture (Kozasa and Gilman, 1996).

### Thermostability Assays (fQCR)

*T*_m_ values were determined using the fast Quantitative Cysteine Reactivity (fQCR) assay as described previously (Isom et al., 2013). Briefly, 10 μL of 40 μM protein was added into 12 strip PCR tubes with 170 μL phosphate buffer pH 7 + 10 μL of 1 mM guanine nucleotide (GDP or GTPγS)+ 10 μL of 500 mM 4-fluoro-7-sulfamoylbenzofurazan (ABDF) for 5 min on ice. The protein was subjected to a temperature gradient in a standard thermocycler for 3 min. The reaction was quenched with ice cold 0.1 N (final concentration) HCl. The ABDF reacts with exposed cysteine residues and emits light detected on a PHERAstar (BMG Labtech) plate reader using wavelengths using excitation and emission bandpass filters of 400 and 500 nm (Isom et al., 2013).

### Nucleotide loading

BODIPYFL-GTPγS (100 nM) was equilibrated in buffer (50 mM HEPES, 10 mM MgCl_2_, 25 mM NaCl, pH 7) for 60 sec. Purified Gα protein (100 nM) was added to a 1 mL cuvette. The fluorescence of the BODIPYFL group (502 nm excitation, 511 nm emission) was measured over 600 sec. 20 μM (final concentration) unlabeled GTPγS was then added to displace BODIPYFL-GTPγS. All measurements were made with a Perkin-Elmer Luminescence Spectrometer and the FLWinLab software package (Jones et al., 2012; McEwen et al., 2001).

### Biolayer Interferometry

Binding between Gα and Gβγ was determined by Biolayer Interferometry (BLI) using Octet Red96 (Fortebio) as described previously (Seneviratne et al., 2011). Briefly, purified biotinylated Gβγ (3 mg/mL) was incubated for 15 min with streptavidin biosensors in either PBST-NGM (25 mM KPO_4_, pH 7, 50 mM NaCl, 0.1% Tween, and 50 μM GDP + 5 mM MgCl_2_) or PBST-NGA (50 μM GDP, 5 mM MgCl_2_, 30 μM AlCl_3_, and 1 mM NaF). Purified Gα_i1_ (untagged) was diluted to 3 μM or 1 μM in PBST-NGM or PBST-NGA and then mixed with Gβγ-loaded sensors for 5 min (association) and then protein-free buffer for 5 min (dissociation) at 25 °C. Nonspecific binding was measured using biosensors that were exposed to buffer alone. Baseline subtraction and Gβγ loading normalization were done in Excel. Kinetic Analysis was done using GraphPad Prism.

### NMR sample preparation and spectroscopy

^15^N-enriched wildtype and R208S Gα_i1_-Δ31 (PET-SUMO-Gα_i1_-Δ31 plasmid) were expressed and purified as described above and as detailed previously (Maly and Crowhurst, 2012). The purified proteins were exchanged into NMR buffer (20 mM sodium phosphate, pH 7.0, 50 mM NaCl, 2 mM MgCl_2_, 200 μM GDP, 5% D2O). To exchange GDP to GTPγS, the proteins were gently exchanged into a GDP-free solution with 1 mM MgCl_2_ and 10-fold excess of GTPγS were added to the protein. The sample was incubated on ice for 30 min and then MgCl_2_ concentration was increased to 5 mM for another 30 min. GTPγS-loaded samples were finally exchanged into NMR buffer with 200 μM GTPγS. Each NMR sample contained 100 μM Gα_i1_-Δ31. NMR spectra were acquired at 25°C on a Bruker Avance 850 NMR spectrometer. Two-dimensional ^1^H–^15^N HSQC experiments were recorded with 1024 and 128 complex points in the direct and indirect dimensions, respectively, 44 scans per increment and a recovery delay of 1.0 sec. Spectral widths used were 13586.957 Hz (^1^H) and 3015.682 (^15^N) Hz. Spectra were processed and analyzed using NMRPipe (NIDDK, NIH) and Sparky (University of California San Francisco). Backbone assignments for wildtype Gα_i1_-Δ31 were transferred from BMRB 30078 for GDP and 26746 for GTPγS (Goricanec et al., 2016). The backbone assignment for Gα_i1_-Δ31-R208S were transferred from the wildype Gα_i1_Δ31. All ambiguously correlated or resolved peaks were omitted during the transfer.

### Starting structural models and the details of molecular dynamics simulations

The initial coordinates of the heterotrimeric stimulatory and inhibitory human G proteins, Gα_s_ and Gα_i1_, were taken from their crystal structures (PDB IDs: 6EG8 and 1GP2 respectively) in the inactive GDP-bound states (Liu et al., 2019; Wall et al., 1995). The initial coordinates of both GTPγS nucleotide as well as the counterion Mg^2+^ were obtained from the crystal structures of the two G proteins in the active state (PDB IDs: 1AZT and 1GIA for Gα_s_ and Gα_i1_ respectively) (Coleman et al., 1994; Sunahara et al., 1997). For building the GTPγS/Mg^2+^-bound heterotrimeric G proteins of each kind the two crystal structures of opposite states were overlaid and GDP/Mg^2+^ in the inactive states were swapped by the GTPγS/Mg^2+^ in the heterotrimeric states. We removed the N-terminus from both the GDP and the GTPγS bound G proteins since this region was found to be missing in the monomeric active state crystals of both G proteins. The GTPγS bound heterotrimeric G proteins of each kind were simulated to capture the early events of dissociation in the Gα and Gβ subunits upon replacing GDP with GTPγS. The GDP-bound G proteins in their resting (inactive) states were also simulated as controls. We used CHARMM-GUI (Jo et al., 2008) for creating input structures for simulating G proteins (G_s_ and G_i_) in aqueous medium. The nucleotides (GDP and GTPγS) and the counterions (Mg^2+^) were parameterized using the CGENFF (Vanommeslaeghe et al., 2012) force field as implemented in CHARMM. Point mutations either disrupting or fostering the proposed triad network in G-proteins were incorporated using the MAESTRO software in Schrodinger (https://www.schrodinger.com/maestro). Each of these heterotrimeric protein-nucleotide complexes was solvated in explicit TIP3P water molecules in a cubic box (approximate dimension of 11.30 nm X 11.30 nm X 11.30 nm) separately and sodium and chloride counterions were added for maintaining the physiological salt concentration of each system at 150 mM. We used the software GROMACS (Hess et al., 2008) (version 2019.4) in combination with the all-atom CHARMM36 (Brooks et al., 2009) force field for performing MD simulations at 310 K coupled to a temperature bath with a relaxation time of 0.1 ps (Berendsen et al., 1984). Pressure was calculated using molecular virial and held constantly by weak coupling to a pressure bath with a relaxation time of 0.5 ps. Each system was first subjected to a 5000 step steepest descent energy minimization for removing bad contacts (Petrova and Solov’ev, 1997). Then, the systems were heated for 100 ps in steps of ramping up the temperature to 310 °K under constant temperature-volume ensemble (NVT). Equilibrium bond length and geometry of water molecules were constrained using the SHAKE algorithm (Andersen, 1983). We used a time step of 2 fs. The short range electrostatic and van der Waals (VDW) interactions were estimated per time step using a charge group pair list with cut-off radius of 8 Å between the centers of geometry of the charged groups. Long range VDW interactions were calculated using a cut-off of 14 Å and long-range electrostatic interactions were treated using the particle mesh Ewald (PME) method (Darden et al., 1993). Temperature was kept constant by applying the Nose-Hoover thermostat (Evans and Holian, 1985). Parrinello-Rahman barostat (Parrinello and Rahman, 1981) with a pressure relaxation time of 2 ps was used for attaining the desired pressure for all simulations. The simulation trajectories were saved each 200 ps for analysis. The protein, nucleotide and counterion atoms were position restrained using a harmonic force constant of 1000 kJ mol-1 nm-2 during the NVT equilibration stage while the water molecules were allowed to move freely around the protein. The system was further equilibrated using the constant pressure NPT ensemble by reducing the force constant on protein, counterion and nucleotide atoms from 5 kJ mol^−1^ nm^−2^ to zero in a gradual manner for 3 ns each while having the pressure coupling on. We also performed an additional 10 ns of unrestrained simulation before beginning the actual production run. This accounts for a total 25 ns of NPT equilibration prior to the production run. We performed three productions runs each 400 ns long starting from three independent sets of initial velocities for each system. Three independent simulations (each 400 ns long) were performed for each system. Thus, we had 1.2 μs long MD trajectory for each of the wildtype Gα_s_ and Gα_i1_ bound to GDP and GTPγS, four Gα_i1_ mutants (R208S, R208Q, E245K and G203A) and four Gα_s_ mutants (R231S, R231Q, E268K and G226A) bound to GTPγS.

### Calculation of root mean square fluctuations (RMSF)

Per residue RMSF for each Gα_i1_ system was computed using the gmx rmsf utility with -res option as implemented in GROMACS. The per residue RMSFs were inserted as B-factor columns of the representative snapshots of the Gα_s_ubunits of each Gα_i1_ system and shown as cartoons.

### Calculation of interaction energy

Short range electrostatic and van der Waals interactions were estimated per time step using a charge group pair list with cut-off radius of 8 Å between the centers of geometry of the charged groups. The residue pairs of the G proteins engaged in van der Waals or electrostatic interactions for > 40% of the whole simulation were considered as sustained contacts and were identified using the GetContacts script available at the GitHub (https://getcontacts.github.io/). These residues of interest were indexed and tagged as energy groups. The total non-bonded interactions energies (van der Waals + Coulombic) between two protein subunits, Gα and Gβ or between Gα and nucleotide were extracted using the gmx energy module of GROMACS. The energies were calculated for each snapshot and averaged over the last 300 ns in each of the three simulation runs for each system. For calculating the interaction energies between Switch II and Switch III regions we used the following definition of Switch II and Switch III regions. Residues G203 to E216 make up the Switch II region for Gα_i1_ and residues G226 to N239 for Gα_s_. Switch III is comprised of residues E236 to N255 for Gα_i1_ and E259 to N278 for Gα _s_.

### Representative structure calculation from MD simulation trajectories

Representative snapshots from the most occupied conformation cluster for each system were chosen for structure representation in the figures. The conformation clustering of the snapshots in the MD trajectories was done using the RMSD-based clustering method using the GROMACS modules gmx rms and gmx cluster with a 1.5 Å cutoff on the concatenated trajectory from three simulations (a total of 1.2 μs simulation time) for each system. The most representative structure of the most populated cluster was calculated as the frame that has the smallest RMSD to the center of this cluster conformation. Snapshots were rendered using VMD (Humphrey et al., 1996) and PYMOL (https://www.schrodinger.com/pymol).

### Principal component analysis and free energy surface generation

For each system in this study, we merged the three independent MD runs into one concatenated trajectory. We then performed the principal component analysis using the gmx covar module of GROMACS for each system using covariance matrix of the Cα atoms of all of the residues. The gmx sham module of GROMACS was used to compute the probability of the microstates and convert it to free energy. The free energy surface thus generated was projected on the principal component space covered by principal component 1 and 2 (PC1 and PC2). We then clustered the MD snapshots by the values of PC1 and PC2 using the in-house extractcluster.py script in MATLAB and calculated the population of each of these conformational clusters (Figure 4). The population of the most populated cluster 1 was compared across the various systems under consideration for Gα_i1_ (Figure S4, Supplemental Information). The MD snapshots under each conformational cluster were concatenated to obtain trajectories for each of these clusters and then the VMD script helicity.tcl was used to calculate the helicity of the Switch II regions in Gα_i1_. The cumulative fraction of variance in the eigenvalues of the PCs (shown in Figure S4, Supplementary Information) accounts for about 95% of the variance captured by the first ten PCs for each Gα_i1_ system.

## SUPPLEMENTAL FIGURE LEGENDS

**Figure S1.**
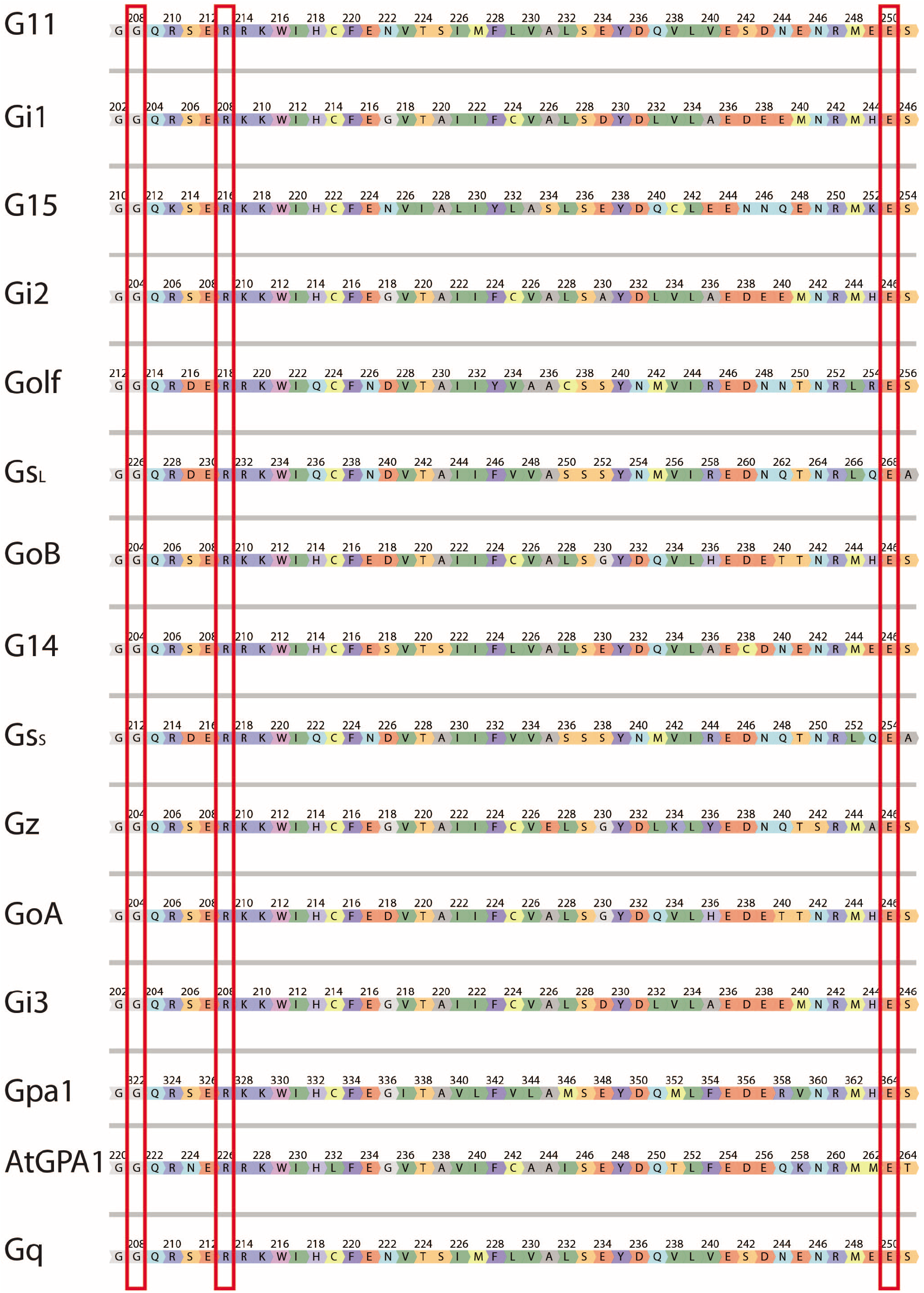
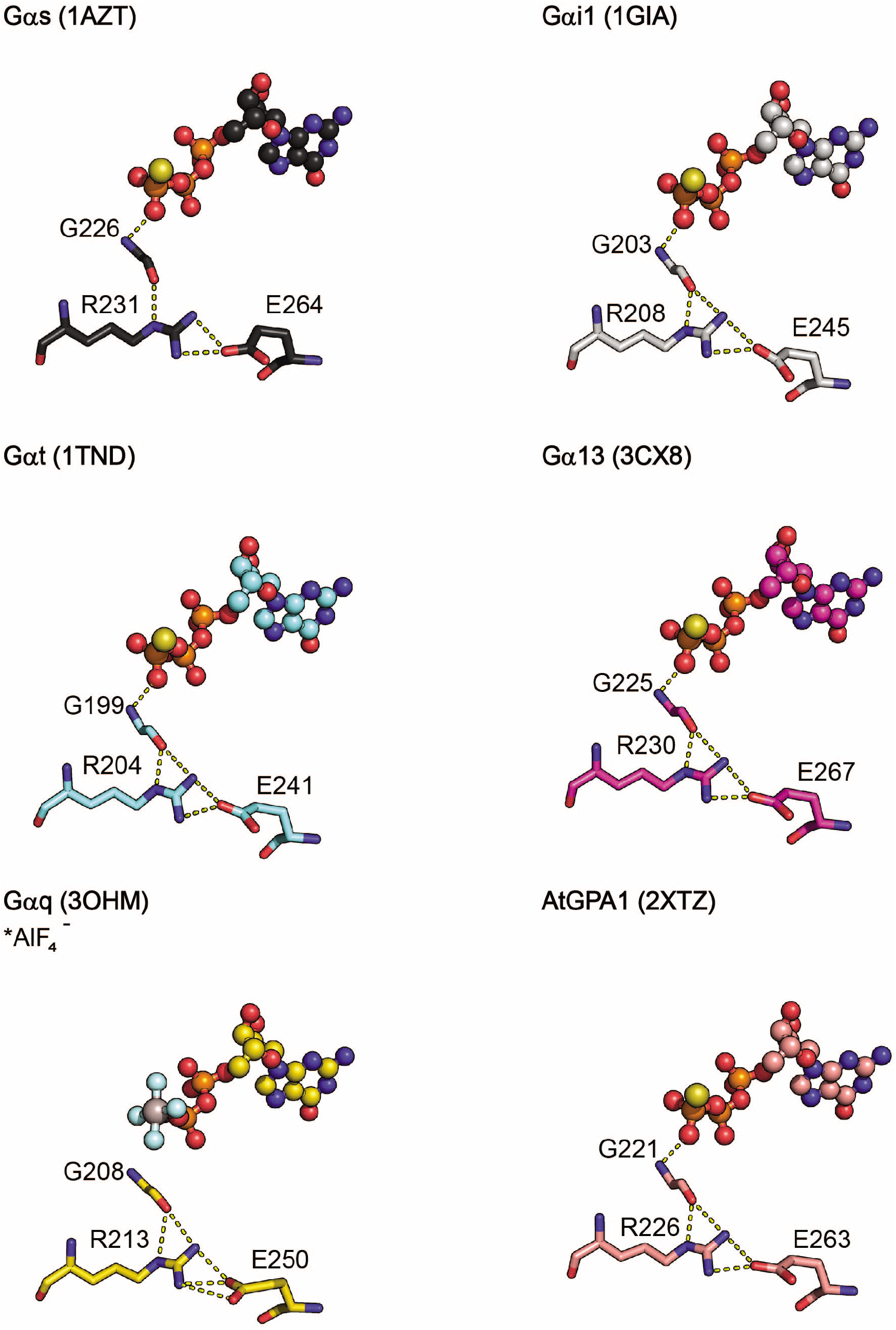
(Related to Figure 1). Multiple sequence alignment. Benchling software was used to align human Gα_s_ubunits as well as Arabidopsis AtGPA1 and yeast Gpa1. Triad residues are indicated by red boxes. Crystal model of Gα_s_, Gα_i1_, Gα_t_ Gα_13_, Gα_q_, AtGPA1 triad residues and GTPγS or GDP-AlF_4_^-^ (ball and stick) from the indicated PDB structures.

**Figure S2 (Related to Figure 2). Linked Excel primer table**

**Figure S3.**
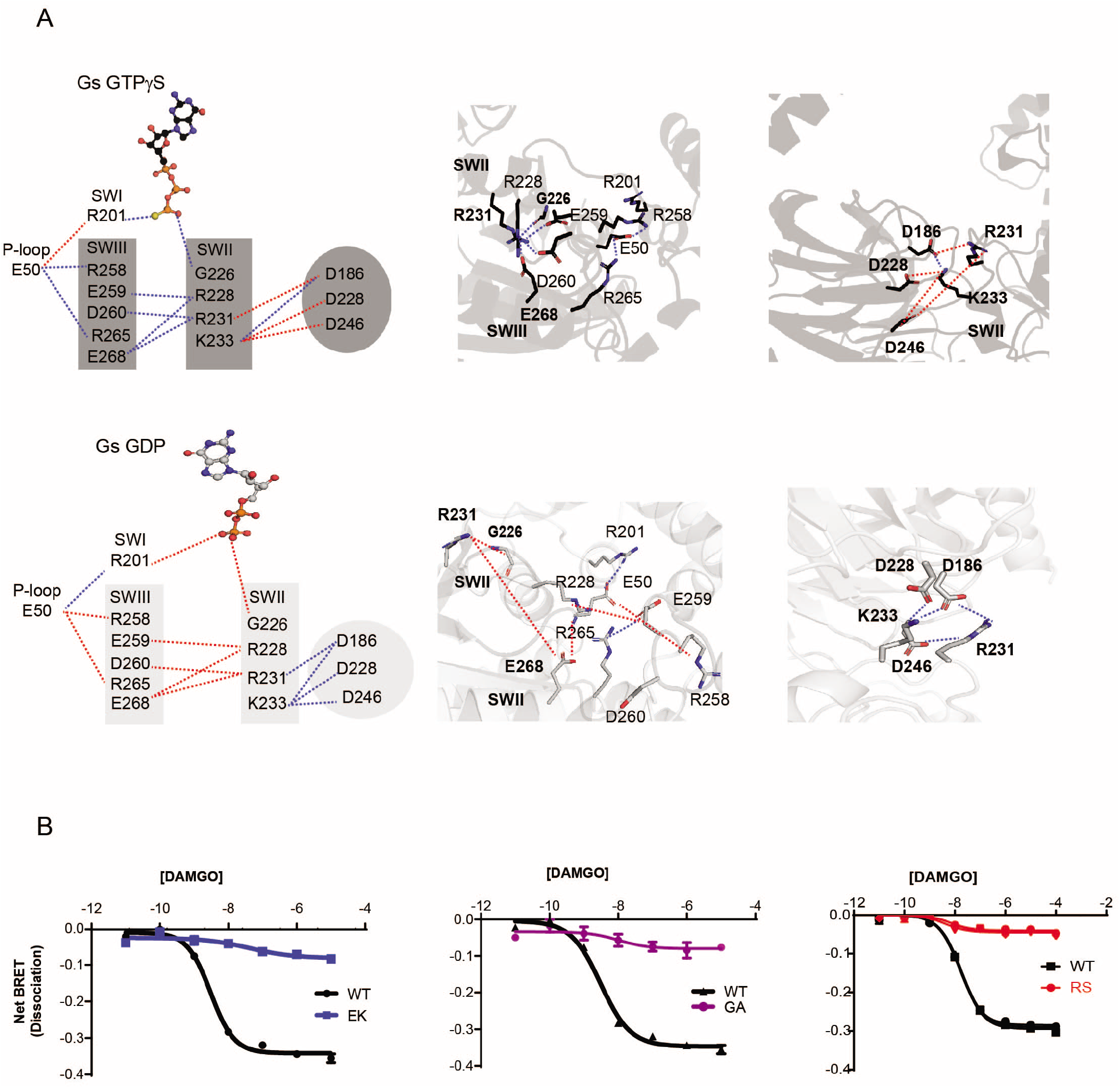
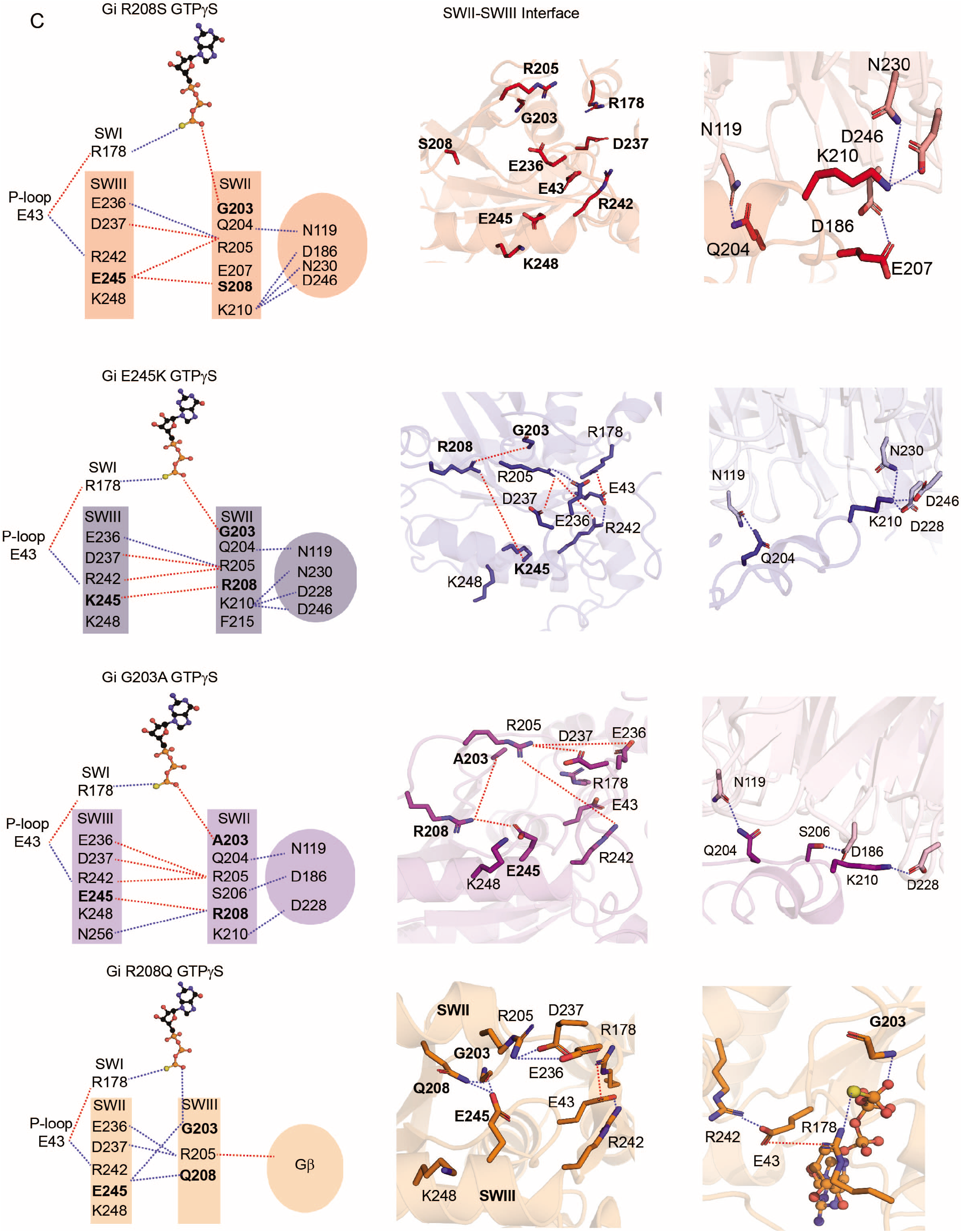
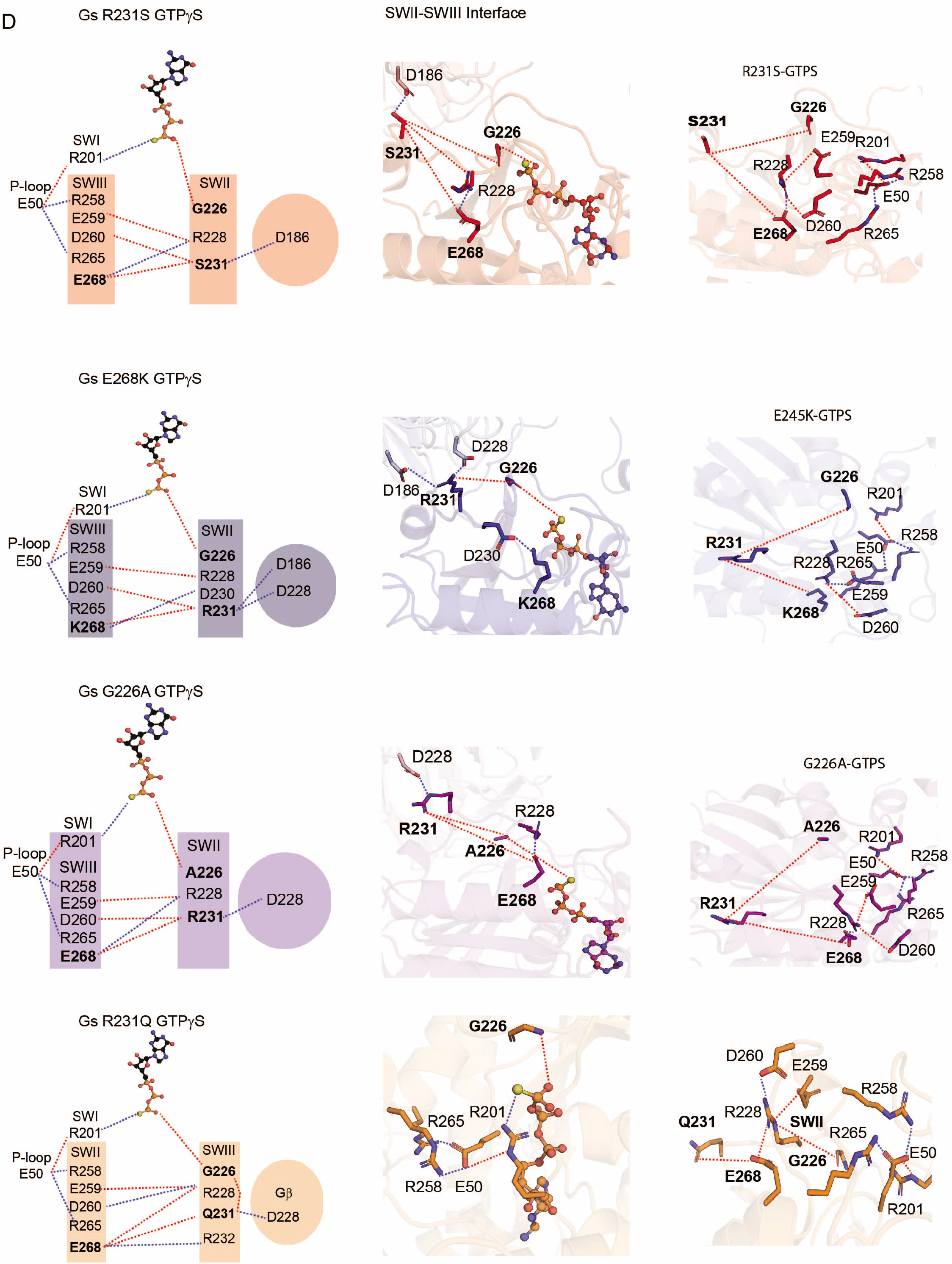
(Related to Figure 3). Gα_s_ network analysis. **(A)** A dynamic mechanism for the effect of the γ-phosphate group triggering the release of the Gβγ subunits from Gα_s_. The dynamic pull-push effect from the γ-phosphate group of GTPγS, that forms the polar residue interaction network involving the P-loop, Switch I, Switch II and Switch III residues, derived from MD simulations for wildtype Gα_s_ bound to GTPγS (top) or GDP (bottom). The nucleotides are shown in ball and stick representation. The blue and the red dotted lines indicate sustained and broken residue contacts, respectively. Right panels show residues involved in the polar network for the GTPγS- and GDP-bound Gα_i1_ switch regions (black and gray colors respectively). **(B**) HEK293 cells transfected with the indicated wildtype or mutant Gα-RLuc8 donor and Gγ-GFP acceptor proteins. Concentration-response measurements (n=2) using μ-opioid receptor are presented as fold-decrease in dynamic range (dissociation, Net BRET). **(C)** A dynamic mechanism for the effect of the γ-phosphate group triggering the release of the Gβγ subunits from Gα_i1_ R208S (red), E245K (blue), G203A (purple), and R208Q (orange). The dynamic pull-push effect from the γ-phosphate group of GTPγS, that forms the polar residue interaction network involving the P-loop, Switch I, Switch II and Switch III residues, derived from MD simulations for wildtype Gα_i1_ bound to GTPγS. **(D)** A dynamic mechanism for the effect of the γ-phosphate group triggering the release of the Gβγ subunits from Gα_s_ R231S (red), E264K (blue), G226A (purple), and R231Q (orange). The dynamic pull-push effect from the γ-phosphate group of GTPγS, that forms the polar residue interaction network involving the P-loop, Switch I, Switch II and Switch III residues, derived from MD simulations for wildtype Gα_s_ bound to GTPγS.

**Figure S4.**
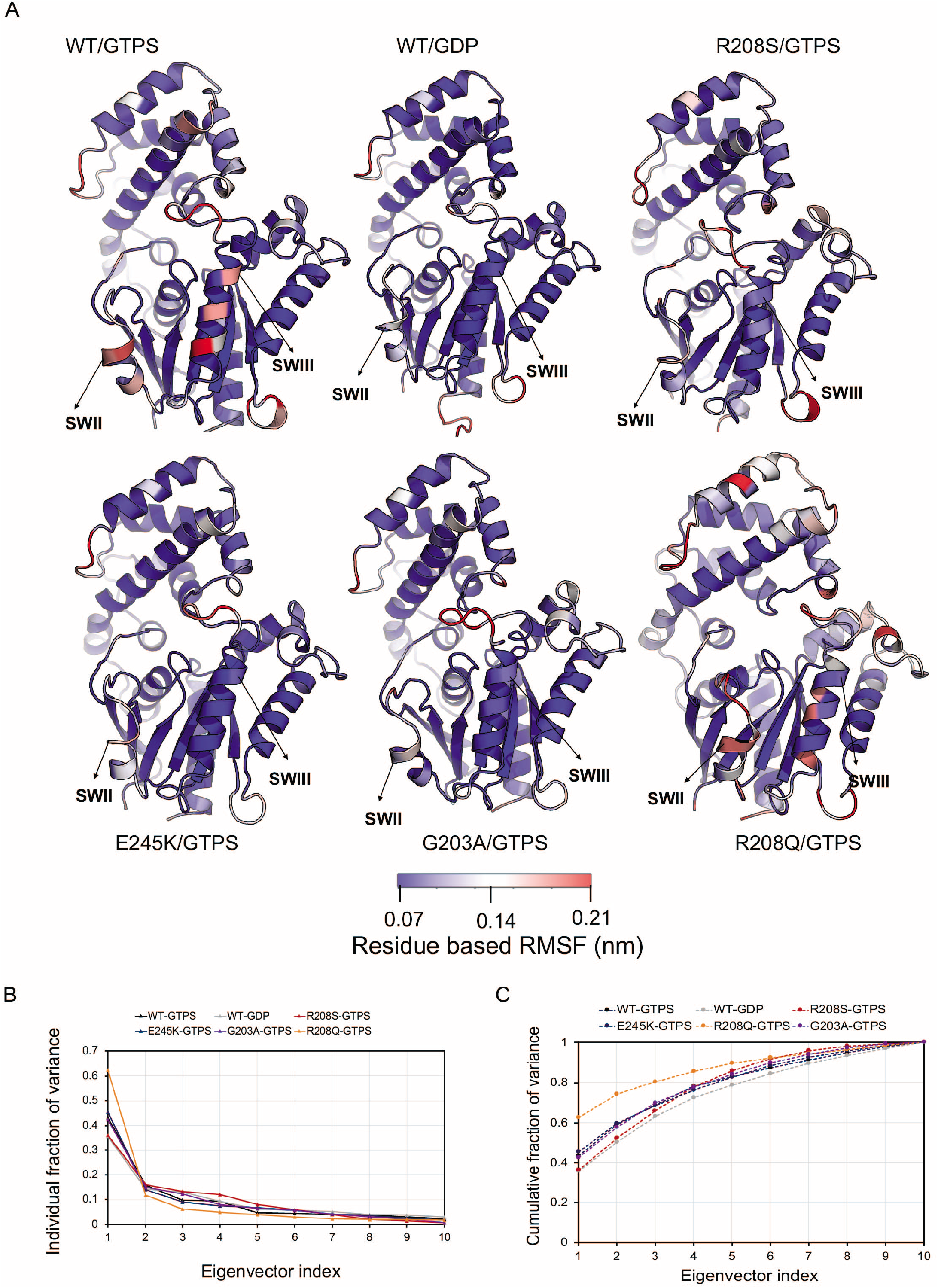
(Related to Figure 4). MD simulations of triad residues that coordinate the active-state conformation. **(A)** and **(B)** Root mean square fluctuations (RMSF) indicating the flexibility of the residues in the protein. The heat map of RMSF is projected on the structures of wild type and various triad mutants of Gα_i_ bound to either GDP or GTP as indicated in the figures. **(C)** The variance in the principal components and its cumulative fraction shown to indicate that the top four principal components carry most of the weight in the principal component analysis that was used to project the free energy surfaces shown in Fig. 4B.

**Figure S5.**
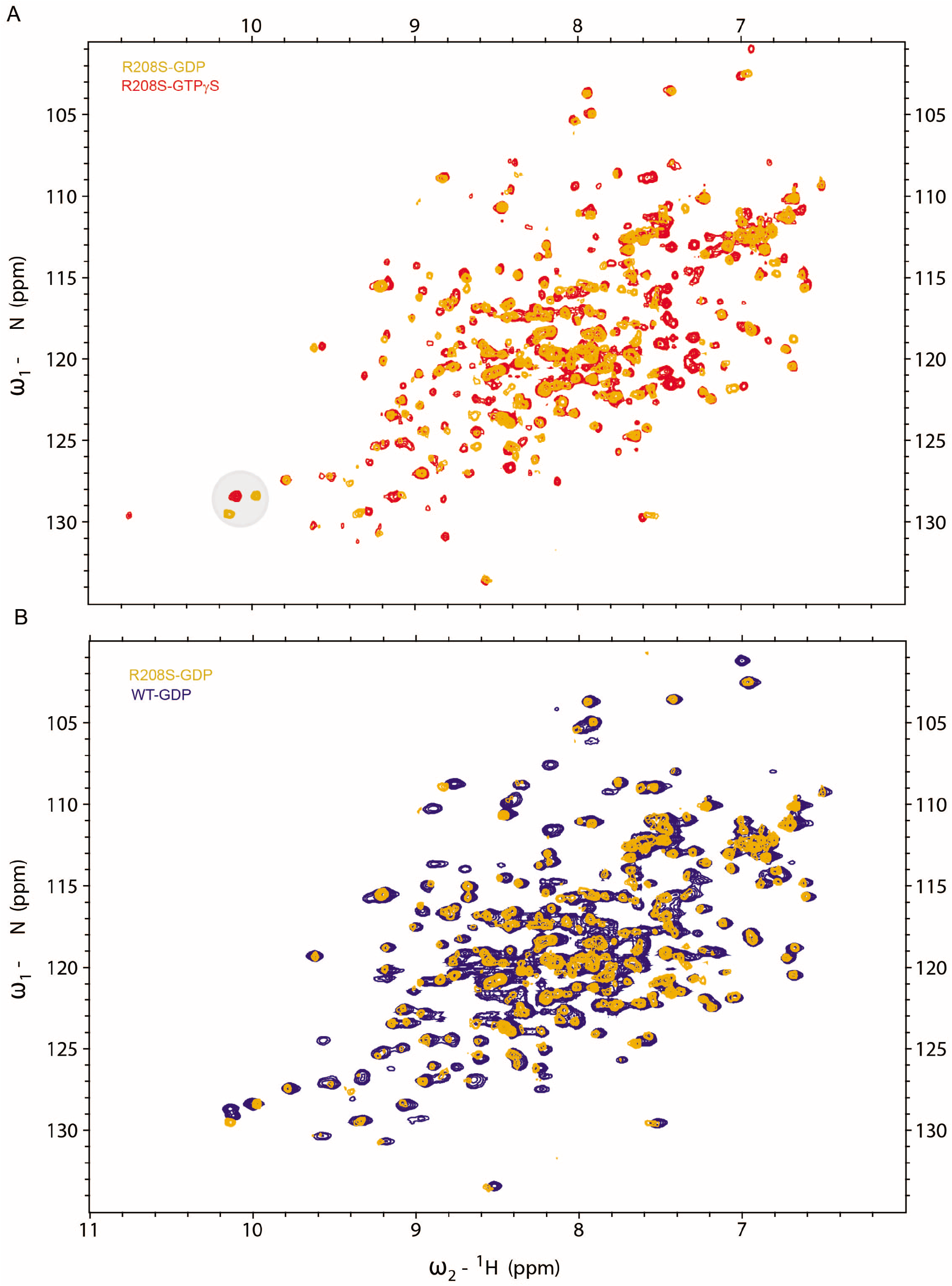
(Related to Figure 5). NMR supplementary spectra. 2D [^15^N, ^1^H]-HSQC NMR of Gα_i1_Δ31 wildtype and R208S mutant. **(A)** Overlay of R208S Gα_i1_ in complex with GTPγS (red), and GDP (orange) **(B)** Overlay of wildtype (blue) and R208S (orange) Gα_i1_ each in complex with GDP

---

**Table S1.**

| $k_{on}$ M<sup>-1</sup> s<sup>-1</sup> | | | | |
| --- | --- | --- | --- | --- |
|  | WT GDP | WT AIF | RS GDP | RS AIF |
| 1 μM | 0.1739 | 0.1089 | 0.1835 | 0.2097 |
| 3 μM | 0.3786 | 0.2191 | 0.3431 | 0.3555 |
| 10 μM | 0.6042 | 0.4996 | 0.5821 | 0.5791 |

| Standard Error in $k_{on}$ | | | | |
| --- | --- | --- | --- | --- |
|  | WT GDP | WT AIF | RS GDP | RS AIF |
| 1 μM | 0.006226 | 0.00811 | 0.005236 | 0.005273 |
| 3 μM | 0.008694 | 0.005218 | 0.007435 | 0.008023 |
| 10 μM | 0.01724 | 0.01252 | 0.01729 | 0.01692 |

| $k_{off}$ s<sup>-1</sup> | | | | |
| --- | --- | --- | --- | --- |
|  | WT GDP | WT AIF | RS GDP | RS AIF |
| 1 μM | 0.1542 | 0.1188 | 0.1532 | 0.1401 |
| 3 μM | 0.2295 | 0.1639 | 0.1634 | 0.1644 |
| 10 μM | 0.2626 | 0.1922 | 0.186 | 0.1786 |

| B max |  |  |  |  |
| --- | --- | --- | --- | --- |
|  | WT GDP | WT AIF | RS GDP | RS AIF |
| 1 μM | 0.1739 | 0.1141 | 0.208 | 0.2525 |
| 3 μM | 0.4217 | 0.3171 | 0.4746 | 0.5472 |
| 10 μM | 0.6234 | 0.6609 | 0.6668 | 0.6999 |

